# Aberrant neuronal cell cycle re-entry induces late-onset Alzheimer’s disease relevant neuropathological and gene expression changes

**DOI:** 10.64898/2026.08.28.747879

**Authors:** Katherine A. Stangis, Ravi S. Pandey, Angela Wang, John S. Beck, Scott E. Counts, Gregory W. Carter, Kevin H.J. Park

## Abstract

Aberrant neuronal cell cycle re-entry (NCCR) is an alternative pathogenic mechanism in Alzheimer’s disease (AD) that has gained substantial support in the literature. The pathogenic role of ectopic NCCR is supported by our past work demonstrating that SV40T-mediated NCCR in adult mice can induce numerous pathologies associated with AD. Since NCCR is chronically induced for an extended period in the mouse model which gives rise to numerous pathologies including neuroinflammation, many of these neuropathological changes could simultaneously participate in driving disease progression. We hypothesized that the NCCR is a primary pathogenic driver and that halting this disease process at a later age could be sufficient for preventing the progression of AD-related pathologies. Here we show that modulation of NCCR at a later age prevents the progression of AD pathologies, including Aβ and tau pathologies. Furthermore, functional genomics analysis demonstrates the late-onset AD (LOAD)-relevance of NCCR. Our findings suggest that our NCCR mouse model could help identify novel therapeutic targets that could aid in preventing AD progression.

## Introduction

Alzheimer’s disease (AD) is the most common cause of dementia, impacting an estimated 50 million individuals worldwide. AD is characterized neuropathologically by amyloid-β (Aβ) plaques, neurofibrillary tangles (NFTs), neuroinflammation and neuronal loss. Much of the current understanding of AD pathogenic mechanisms centers around Aβ plaques and the amyloid hypothesis, which is based in large part on studies utilizing familial AD mouse models that express amyloid precursor protein (APP) and presenilin gene mutations that represent 1-2% of AD cases (Jankowsky and Zheng, 2017). These mice have been essential in elucidating the pathogenic role of Aβ peptides while suggesting that Aβ alone is insufficient for inducing neurodegeneration since transgenic overexpression of mutant APP and mutant presenilin in mice results in robust generation of Aβ peptides and plaques, but do not display NFT-related tau pathology. Additionally, the majority (90-95%) of the AD cases are sporadic, and not much is known about non-genetic pathogenic processes that drive many of the AD-related pathologies.

Aberrant neuronal cell cycle re-entry (NCCR) is an alternative pathogenic mechanism in AD that has gained substantial support in the literature. Ectopic neuronal cell cycle activation as a contributing factor in AD pathogenesis is suggested by the presence of various cell cycle markers in familial and sporadic AD postmortem brains (Arendt et al., 2010; Boeras et al., 2008; Busser et al., 1998; Herrup and Yang, 2007; Kruman et al., 2004; McShea et al., 1997; Nagy et al., 1997; Vincent et al., 1996; Yang et al., 2003, 2001). Between 5 and 10% of the cells in the hippocampus, cortex, locus coeruleus, nucleus basalis, and dorsal raphe nucleus show cell cycle marker labeling in post-mortem brain samples from mild cognitive impairment and AD subjects, whereas no significant expression of cell cycle activation markers is observed in control brains (Busser et al., 1998; Yang et al., 2003). A network analysis using epigenetic and transcriptomic datasets from AD brains identified major hub genes associated with molecular pathways involved in cell cycle re-entry and inflammation (Li et al., 2019). Furthermore, forced cell cycle re-entry in postmitotic neurons leads to neurodegeneration and cell death (Herrup et al., 2004). Thus, several lines of evidence support a role for aberrant cell cycle re-entry as a pathogenic factor in AD.

The pathogenic role of ectopic NCCR is supported by our work demonstrating that SV40T-mediated NCCR in adult mice can induce numerous pathological hallmarks associated with AD (Barrett et al., 2021; Park et al., 2007; Park and Barrett, 2020). Moreover, our recent findings showed that NCCR enhances and generates many of the neuropathological features associated with AD in *App^NL-F^* knock-in (KI) mice, namely PHF-1-positive phospho-tau lesions, neuroinflammation, brain leukocyte infiltration, DNA damage responses, and cortical atrophy (Barrett et al., 2021). However, the Aβ plaque load was not altered in the NCCR-*App^NL-F^* mice at 9, 12, or 18 months of age (Barrett et al., 2021). We postulate that the effect of cell cycle re-entry on Aβ plaque deposition was not observable due to the excessive levels of Aβ42 produced in NCCR-*App^NL-F^* mice resulting from the presence of both Swedish and Iberian mutations (Saito et al., 2014).

To evaluate the effect of NCCR on Aβ42 production in humanized Aβ *App* KI mice, we utilized the *App^NL^* knock-in mice in combination with our NCCR mouse model (Barrett et al., 2021; Park et al., 2007; Park and Barrett, 2020). *App^NL^* mice have a humanized Aβ domain and possess the Swedish mutation only (Saito et al., 2014). The gene construct in *App^NL^* mice is similar to *App^NL-F^* mice in all aspects except for lacking the Iberian mutation. However, the notable difference between the two models is that, unlike the *App^NL-F^* mice, *App^NL^* mice are characterized by preferential production of Aβ40, but do not display plaques, neurodegeneration, or neuroinflammation at any age (Masuda et al., 2016; Saito et al., 2014; Salas et al., 2018). In addition, we evaluated the differential gene expression patterns of 18-month-old NCCR-*App^NL^* mice compared to 18-month-old *App^NL^* mice using the NanoString nCounter AD panel. This gene-set panel was developed to correlate human disease processes and pathways with mouse brain transcripts as a means for evaluating the relevance of familial AD mouse models for studying LOAD (Preuss et al., 2020; Wan et al., 2020). We also evaluated SV40T-positive neurons in 4-month-old NCCR-AD mice using laser capture microdissection-microarray (LCM-microarray) analysis.

We demonstrate that, in contrast to *App^NL^* mice, the NCCR-*App^NL^* mice display intraneuronal Aβ42, neuroinflammation, brain leukocyte infiltration, and neurodegeneration. However, Aβ plaque deposits were not detected at either 12 or 18 months of age. Interestingly, PHF-1-positive phospho-tau lesions were also not observed in these animals. Gene expression analysis of 18-month old NCCR-*App^NL^* and *App^NL^* mice using the NanoString mouse AD panel identified 223 differentially expressed genes (FDR adjusted p<0.05) that show statistically significant correlation with AD co-expression modules for various functional consensus clusters. Furthermore, silencing of neuronal expression of SV40T starting at 12 months of age (i.e., after 11 months of NCCR) arrested the progression of AD-associated neuropathological features observed at 18 months of age, further illustrating the potential pathogenic role of NCCR in AD. Overall, our findings demonstrate that ectopic NCCR can give rise to diverse neuropathological and gene expression changes linked with LOAD. Furthermore, these results highlight the relevance and usefulness of our NCCR mice as a LOAD mouse model for investigating the complex interaction between AD-associated pathologies, and identifying new therapeutic targets and compounds.

## Materials and Methods

### Animals

All mice used for this study were maintained on a C57BL/6N background. *App^NL^* and *App^NL-F^* and mice were supplied by the Saido Laboratory and have been described previously (Saito et al., 2014). Camk2a-tTA mice were obtained from the Jackson Laboratory (Bar Harbor, ME) and maintained as an in-house breeding line. TRE-SV40T mice, which have been previously generated and characterized, were similarly maintained in-house (Park et al., 2007)(Barrett et al., 2021). *App^NL^* mice were crossed with both Camk2a-tTA and TRE-SV40T mice to generate Camk2a-tTA^+/-^/*App^NL/NL^* and TRE-SV40T^+/-^/*App^NL//NL^* lines. The same was done with *App^NL-F^* mice to generate Camk2a-tTA^+/-^/*App^NL-F/NL-F^* and TRE-SV40T^+/-^/ *App^NL-F//NL-F^* lines. Camk2a-tTA^+/-^ /*App^NL/NL^* mice were then crossed with TRE-SV40T^+/-^/ *App^NL//NL^* mice to produce Camk2a-tTA^+/-^/TRE-SV40T^+/-^/*App^NL//NL^* mice (hereby known as the NCCR-*App^NL^* mouse model). The same was done for the *App^NL-F^* lines to produce Camk2a-tTA^+/-^/TRE-SV40T^+/-^/*App^NL-F//NL-F^* mice (hereby known as the NCCR-*App^NL-F^* mouse model). Breeding pairs were maintained on a doxycycline (dox) diet to prevent SV40T expression within offspring *in utero,* and until the time of weaning. Offspring were weaned at 21 days of age and maintained on dox diet until reaching one month of age. At this point, offspring were switched to a standard diet to induce neuronal cell cycle re-entry (NCCR). The mice were continuously maintained on dox diet until 12 or 18 months of age for evaluation.

For halting the NCCR starting at 12 months of age, a cohort of mice that were maintained on standard diet were put back on dox diet at 12 months of age and continuously maintained on dox diet until 18 monhths of age for evaluation. A cohort of *App^NL^* and *App^NL-F^* were also put back on dox diet as diet control groups. All animals were maintained on a 12-hour light cycle and group housed in individually ventilated cages with food and water available *ad libitum.* Tail tissue was collected from all animals at the time of weaning and again at the animal’s endpoint for genotyping PCR. All animal protocols were approved by the Institutional Animal Care and Use Committee at Central Michigan University.

### Antibodies

The following primary antibodies were utilized for this study: SV40T and PCNA (PC10 and Pab 101, respectively, Santa Cruz Biotechnology, Dallas, TX, USA); PHF-1 antibody (generously provided to us by Dr. Peter Davies, The Feinstein Institute for Medical Research, Manhasset, NY); 6E10 (Biolegend, San Diego, CA, USA); C42 (C-terminal human Aβ_42_, IBL-America, Minneapolis, MN); C40 (C-terminal human Aβ_40_, IBL-America, Minneapolis, MN); GFAP (MAB360, Millipore, Billerica, MA, USA); Iba1 (178846, Abcam, Cambridge, MA, USA); Hoechst (Thermo Fisher Scientific, Waltham, MA, USA), and NeuN (MAB377, Millipore, Billerica, MA, USA). Secondary Antibodies utilized: Alexa-fluorine conjugated goat-anti rabbit 488, goat-anti rabbit 594, goat-anti mouse 594, and goat-anti rat 594 (Thermo Fisher Scientific, Waltham, MA, USA).

### Immunofluorescence

Immunofluorescence staining was performed as was previously described (Barrett et al., 2019). In brief, mice were anestitized using isofluorine, perfused using a saline solution followed by 4% paraformaldehyde, and whole brains were retrieved. Tissue was stored in 4% paraformaldehyde for 24 hours, and then stored in phosphate buffered saline (1xPBS) until sectioning. Coronal sections, 30 microns thick, were cut using a vibratome (Leica VT1000 S, Leica Biosystems, Buffalo Grove, IL, USA). Sections were stored in a cryoprotective buffer at -20°C until time of staining. Staining involved numerous rinses, permeabilization, and blocking followed by overnight primary antibody incubation at 4°C. Sections were then rinsed again the following day and incubated in the appropriate fluorescent secondary antibodies as previously listed. Following a rinse step, sections were briefly incubated in Hoechst (1:5000 in 1X tris buffered saline (1X TBS)), rinsed, and finally mounted on Superfrost Plus microscope slides (Thermo Fisher Scientific, Waltham, MA) and cover-slipped using Fluromount G (Thermo Fisher Scientific, Waltham, MA). Certain antibodies required antigen retrieval prior to the protocol described above. When staining using C40 and C42, sections were briefly incubated in 90% formic acid for 5 minutes, then rinsed in distilled water for 5 minutes before proceeding with the staining procedure described above. When staining using SV40T and PCNA, sections were incubated in 10mM Sodium Citrate (pH 6.0), floating in a 95°C water bath for 10 minutes, then allowed to cool at room temperature for 20 minutes before proceeding with the staining procedure described above. Images were taken utilizing a Zeiss AxioCam M2 microscope (Carl Zeiss Inc., Thornwood, NY) and digitized using ZEN 2.6, Blue Edition software (Carl Zeiss Inc., Thornwood, NY).

### ImageJ Analysis

Cortical area fraction covered by GFAP signal was quantified using ImageJ. Analysis was performed on coronal sections containing cortical and hippocampal regions that were matched across brain samples, which were region matched to the rostral two sections utilized for quantifying Aβ protein fragment load. Digitized images were converted to 16-bit greyscale, and threshold was adjusted appropriately to minimize background and artifact. Threshold intensity was adjusted similarly for all sections across ages and groups. The whole cortex of each section was traced, and the area fraction of the total cortex occupied by GFAP signal was quantified.

### Stereological Analysis

Cell quantification and cortical area quantification was conducted through the use of the Optical Fractionator Probe within the Stereo Investigator software (MicroBrightField, Williston, VT,USA). To quantify the proportion of neuronal cells expressing PCNA, PCNA/C42, or C40, one region matched coronal section containing cortical and hippocampal regions per animal was utilized. To quantify CD45^+^/Iba1^-^, CD45^+^/Iba1^+^, or C42 cells, three region matched coronal sections across brain samples were assessed. The cortical area was assessed using the following parameters: for PCNA/C42, C40 and C42 quantification: 400×400μm counting frame, 800×800μm sampling grid. The cortical and hippocampal areas were assessed using the following parameters for CD45^+^/Iba1^-^ and CD45^+^/Iba1^+^ cell quantification: 400×400μm counting frame, 800×800μm sampling grid. Each section analyzed was used as an individual data point, as previously reported (Barrett et al., 2021, 2019). To quantify cortical thinning, five coronal brain sections per animal were utilized. These coronal brain sections were region matched across brain samples. The whole cortical area of each section was traced and quantified.

### Statistical analysis of neuropathological assessments

All neuropathological data were assessed using two-way ANOVA with Tukey’s post-hoc test. For sex difference assessment, male and female data within genotype group were independently evaluated (sex as main effect) via two-way ANOVA with Tukey’s post-hoc test (age x sex). When Tukey’s post-hoc test did not show male vs female difference within a genotype at a given age, the data from males and females were combined for two-way ANOVA analysis (genotype x age). Two-way ANOVA results are provided in Excel files as Supplementary Tables 1-3.

### Gene expression analysis using NanoString gene mouse AD Panel

The NanoString Mouse AD gene expression panel (Preuss et al., 2020) was used for gene expression profiling on the nCounter platform (Geiss et al., 2008) (NanoString, Seattle, WA). Mouse NanoString data were collected from brain hemispheres from 18-months-old *App^NL^* (n=6; 2M, 4F) and NCCR-*App^NL^* (n=6; 2M, 4F) mice. nSolver software was used for generating NanoString gene expression count data for these mice. Normalization was done by dividing counts within a lane by geometric mean of the housekeeping genes from the same lane (Preuss et al., 2020). Next, normalized count values were log-transformed for downstream analysis. Differential gene expression analysis for NCCR-*App^NL^* mice compared to the *App^NL^* mice was performed using the voom-limma package in R (Ritchie et al., 2015).

### Human post-mortem brain cohorts and gene co-expression modules

Data on the 30 human brain co-expression modules (Wan et al., 2020) based on meta-analysis of differential gene expression from seven brain regions in postmortem samples obtained from three independent LOAD cohorts (Allen et al., 2016; De Jager et al., 2018; Wang et al., 2018) was obtained from the Synapse data repository (https://www.synapse.org/#!Synapse:syn11932957/tables/; SynapseID: syn11932957). These 30 human brain co-expression modules were further grouped into five consensus clusters that describe the major functional groups of alterations observed in human AD (Preuss et al., 2020; Wan et al., 2020). A detailed description on how co-expression modules were identified can be found in the recent study that identified the harmonized human co-expression modules as part of transcriptome wide AD meta-analysis (Wan et al., 2020).

### Mouse-human expression comparison

Next, we assessed the effect of sex and NCCR (neuronal cell cycle re-entry) by fitting a multiple regression model using the lm function in R as (Pandey et al., 2019):

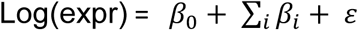

Where the sum is over sex (male) and NCCR. The log(expr) represents the log-transformed normalized count of each gene in the NanoString gene expression panel. To compare mouse expression changes with those observed in human disease, we computed Pearson correlations between gene expression changes (log fold change) in human AD cases versus controls and the effect of sex and NCCR as measured above for each gene in each human brain co-expression module (Greenwood et al., 2020; Wan et al., 2020) using cor.test function built in R as:

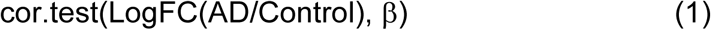

from which we obtained both the correlation coefficient and the significance level (p-value) of the module-level correlation. Log_2_FC values for human transcripts were obtained through the AD Knowledge Portal (Greenwood et al., 2020) (https://www.synapse.org/#!Synapse:syn14237651).

### LCM-microarray analysis

Tissue sections from four-month-old NCCR mice (n = 4) containing cortical and hippocampal regions were immunostained using a mouse monoclonal antibody to SV40T-antigen (clone PAB101; 1:50; Santa Cruz Biotechnology, Dallas, TX). Unlabeled (SV40T-) and labeled (SV40T+) neurons (n = 200 neurons/group) were accessed using a Leica LMD6500 LCM instrument (Leica Biosystems, Deer Park, IL) and analyzed by the custom microarrays (Counts et al., 2014, 2007; Ginsberg et al., 2010; Tiernan et al., 2018, 2016). Amplification of SV40T+ and SV40T-neuronal RNA was performed using terminal continuation (TC) methodology (Counts et al., 2014)(Counts et al., 2007)(Ginsberg et al., 2010)(Tiernan et al., 2016)(Tiernan et al., 2018). Briefly, LCM-accessed neurons (n = 50/sample, 4 samples/group) were homogenized in 500 μL Trizol reagent (Invitrogen, Carlsbad, CA). RNAs were reverse transcribed in the presence of the poly d(T) primer (100 ng/μl) and TC primer (100 ng/μl) in 1× first strand buffer (Life Technologies), 2 μg of linear acrylamide (Applied Biosystems), 10 mM dNTPs, 100 μM DTT, 20 U of SuperRNase Inhibitor (Life Technologies), and 200 U of reverse transcriptase (Superscript III, Life Technologies). Single-stranded cDNAs were digested with RNase H and re-annealed with the primers in a thermal cycler: RNase H digestion step at 37 °C, 30 minutes; denaturation step 95 °C, 3 minutes; primer re-annealing step 60 °C, 5 minutes. This step generated cDNAs with double-stranded regions at the primer interface. Samples were then purified by column filtration (Montage PCR filters; Millipore, Billerica, MA), divided into two technical replicates, and hybridization probes were synthesized by *in vitro* transcription using ^33^P incorporation in 40 mM Tris (pH 7.5), 6 mM MgCl2, 10 mM NaCl, 2 mM spermidine, 10 mM DTT, 2.5 mM ATP, GTP and CTP, 100 μM of cold UTP, 20 U of SuperRNase Inhibitor, 2 KU of T7 RNA polymerase (Epicentre, Madison, WI), and 120 μCi of 33P-UTP (Perkin-Elmer, Boston, MA) (Counts et al., 2014, 2007; Ginsberg et al., 2010; Tiernan et al., 2018, 2016). The reaction was performed at 37 °C for 4 h. Radiolabeled TC RNA probes were hybridized to custom-designed microarrays without further purification. Arrays were hybridized overnight at 42 °C in a rotisserie oven and washed sequentially in 2X SSC/0.1% SDS, 1X SSC/0.1% SDS, and 0.5X SSC/0.1% SDS for 20 min each at 42 °C. Arrays were placed in a phosphor screen for 24 h and developed on a Bio-rad Molecular Imager (Hercules, CA).

### Custom-designed microarray platforms and data collection

Array platforms consisted of 1 μg linearized cDNA purified from plasmid preparations adhered to high-density nitrocellulose (Hybond WL, GE Healthcare). Approximately 864 cDNAs of interest to neurobiology were utilized on the array platform. Hybridization signal intensity was determined using Bio-Rad Image Lab software. Expression of TC amplified RNA bound to each linearized cDNA corrected for background signal was expressed as a ratio of the total hybridization signal intensity of the array (i.e., global normalization) (Counts et al., 2014, 2007; Ginsberg et al., 2010; Tiernan et al., 2018, 2016). The data analysis generated expression profiles of relative changes in mRNA levels among the phenotypically distinct SV40T+ and SV40T-neurons.

### Pathway and Gene set enrichment analysis

KEGG pathway and Gene Ontology (GO) enrichment analyses were performed using clusterProfiler package within the R software environment (Yu et al., 2012). The enrichKEGG and enrichGO functions were used for enrichment of KEGG pathways and Gene Ontology biological processes, respectively. Pathways were determined to be significant after multiple testing correction (FDR adjusted p < 0.05). Gene set enrichment analysis (GSEA) (Subramanian et al., 2005) was performed to identify significantly up and down-regulated gene sets for each genetic risk variant as implemented in the clusterProfiler package for the KEGG and REACTOME pathway library. NanoString AD panel genes were ranked based on regression coefficient calculated for each mouse variants. Enrichment scores for all associated KEGG and REACTOME pathways were computed to compare relative expression on the pathway level between NCCR, 5XFAD, APOE4, and Trem2.R47H and other LOAD mouse models (Preuss et al., 2020).

## Results

### Neuronal cell cycle re-entry induces intraneuronal Aβ42 accumulation in App^NL^ mice but not plaques

*We crossed App^NL^* mice with our NCCR mice in order to examine the effect of ectopic neuronal cell cycle re-entry in *App* knock-in mice possessing humanized Aβ domain, but lacking AD-related pathologies (Masuda et al., 2016; Saito et al., 2014; Salas et al., 2018). *App^NL^* animals predominantly generate Aβ_40_ (Saito et al., 2014), therefore we tested to see whether aberrant neuronal cell cycle re-entry could produce Aβ_42_ and plaques in these animals using either C-terminal Aβ_42_ (C42) or C-terminal Aβ_40_ (C40) antibody to distinguish Aβ(x-42) species from Aβ(x-40) species. Intracellular Aβ(x-40) immunoreactivity was observed in both NCCR-*App^NL^* and *App^NL^* mice at 12 and 18 months of age (Supplementary Figure 1A). On the other hand, intraneuronal Aβ(x-42) is detected only in the 12- and 18-month old NCCR-*App^NL^* animals, whereas age-matched *App^NL^* animals did not display any C42 immunoreactivity (Supplementary Figure 1A). There was no genotype or sex difference in the number of either Aβ(x-40) or Aβ(x-42) immunopositive cells (Supplementary Figure 1B,C; Supplementary Table 1). The number of cells with intraneuronal Aβ(x-42) labeling was 111% higher in 18-month old *NCCR-App^NL^* animals compared to 12-month old *NCCR-App^NL^* animals demonstrating age-dependent increase (Figure 1A). On the other hand, the number of Aβ(x-40) bearing cells did not show either NCCR- or age-dependent changes (Figure 1B). Despite the presence of intraneuronal Aβ(x-42), we did not detect any Aβ plaques in the animals with either C42 or 6E10 antibodies (Supplementary Figure 1D). We also did not observe any notable PHF-1+ phospho-tau lesions in the NCCR-*App^NL^* mice (Supplementary Figure 1E).

**Figure 1.**
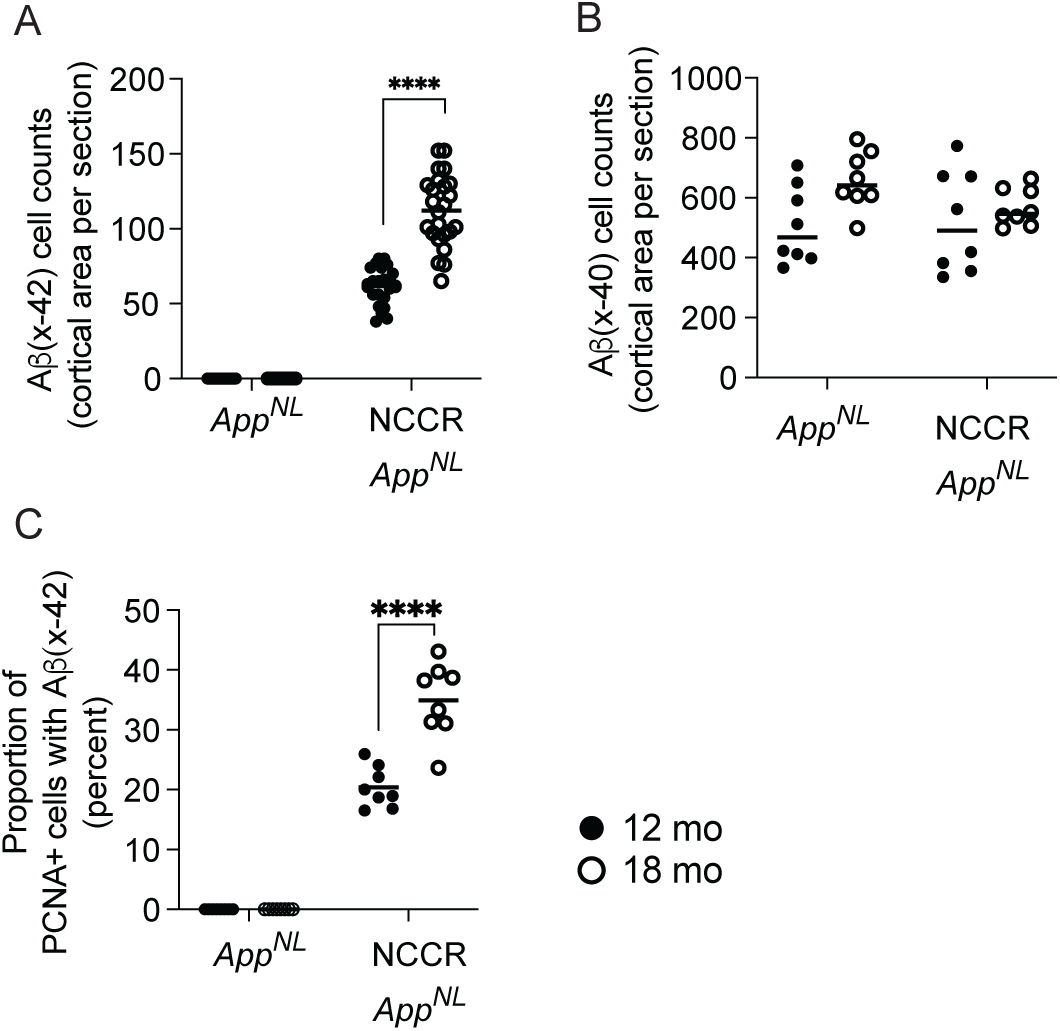
NCCR promotes intraneuronal Aβ(x-42) in *App^NL^* mice. Aβ(x-42) was labeled using C-terminal Aβ42 antibody in coronal sections from 12- and 18-month old mice (Supplementary Figure 1). (A) intraneuronal Aβ(x-42) labeling was detected in NCCR-*App^NL^* brain sections but was absent in *App^NL^* brain sections. Comparison of number of cells bearing intraneuronal Aβ(x-42) labeling in the cortical area between 12- and 18-month old NCCR-*App^NL^* animals show significant increase with age (upaired t-test, p<0.0001). (B) All cells bearing intraneuronal Aβ(x-42) labeling were co-labeled for PCNA. However, only a proportion of PCNA labeled neurons were co-labeled for Aβ(x-42), which increased with age (12mo vs 18mo NCCR-*App^NL^*, unpaired t-test, p<0.0001). (C) Aβ(x-40) was labeled using C-terminal Aβ40 antibody in coronal sections. Number of cells bearing Aβ (x-40) labeling was not affected by neuronal cell cycle re-entry, however, there was a significant age effect (two-way ANOVA: p=0.0276). Complete two-way ANOVA results are provided in suppl. table 2. The following numbers of animals were examined: at 12 mo: NCCR-*App^NL^* (n=4M,4F); *App^NL^* (n=4M,4F); at 18 mo: NCCR-*App^NL^* (n=4M,4F); *App^NL^* (n=4M,4F). The numbers of matched coronal sections measured across the animals for the following labeling: Aβ(x-42), n=3; Aβ(x-40), n=1. Each data point represents a measure from each section.

As anticipated, the neuronal expression of SV40T was observed only in NCCR-*App^NL^* mice. Additionally, neuronal cell cycle re-entry in animals expressing SV40T was confirmed through the use of proliferating cell nuclear antigen (PCNA) antibody, and PCNA immunoreactivity was detected only in NCCR-*App^NL^* animals. Notably, dual-label immunofluorescence using PCNA and C42 antibodies showed co-labeling and demonstrated intraneuronal Aβ(x-42) accumulation that is restricted to cells with PCNA expression. However, only a proportion of PCNA+ cells in both 12- and 18-month old NCCR-*App^NL^* mice showed co-labeling with C42 (Fig 1C), which did not show a sex effect (Supplementary Figure 2A; Supplementary Table 1). The proportion of PCNA+ cells with intraneuronal Aβ(x-42) accumulation was 32 percent higher in the 18-month old compared to 12-month old mice (Figure 1C). However, the number of PCNA expressing neurons was not different between 12- and 18-month old NCCR-*App^NL^* animals (Supplementary Figure 2B). Evaluation of Hoechst DNA signal in SV40T+ and PCNA+ neurons showed statistically significant differences in the intensity by both immunopositivity and age (Supplementary Figure 2C; Supplementary Table 2). Post-hoc analysis showed significant increase in the Hoechst DNA signal in the 18-month-old NCCR-*AppNL* animals compared to the other groups (Supplementary Figure 2C). These data suggest increased DNA synthesis in the NCCR animals, which is increased in older NCCR mice.

### Neuronal cell cycle re-entry (NCCR) induces neuroinflammation and cortical atrophy in App^NL^ mice

Although *App^NL^* mice possess a humanized Aβ domain bearing the Swedish familial AD mutation, these mice do not display any AD-associated pathologies (Saito et al., 2014). We demonstrated previously that the induction of neuronal cell cycle re-entry enhances the activation of microglia and astrocytes in wild-type and *App^NL-F^* knock-in mice (Barrett et al., 2021; Park and Barrett, 2020). Therefore, we evaluated neuroinflammation as well as brain leukocyte infiltration and cortical atrophy in 12- and 18-month old NCCR-*App^NL^* animals to determine whether NCCR can induce these changes in the *App^NL^* animals.

In agreement with our previous findings (Barrett et al., 2021), NCCR increased the number of activated microglia (Figure 2A) and brain infiltrating leukocytes (Figure 2B) in both the cortex and hippocampus, with a clear age-dependent increase for both measures in both the cortex and hippocampus of *App^NL^* mice. There was no sex difference detected for either activated microglia (Supplementary figure 3A; Supplementary Table 1) or brain infiltrating leukocytes (Supplementary Figure 3B; Supplementary Table 1). By contrast, both male and female NCCR-*App^NL^* animals demonstrated an increase in astrocytosis at 12- and 18-months of age compared to *App^NL^* animals (Figure 2C) with an observed sex difference favoring higher cortical GFAP levels in females (Supplementary Figure 3C; Supplementary Table 1). In addition, there was a statistically significant age-dependent GFAP increase in male NCCR-*App^NL^* animals (Figure 2C). However, this was not the case for the female NCCR-*App^NL^* animals suggesting a possible sex difference in astrocyte response to NCCR with age (Figure 2C).

**Figure 2.**
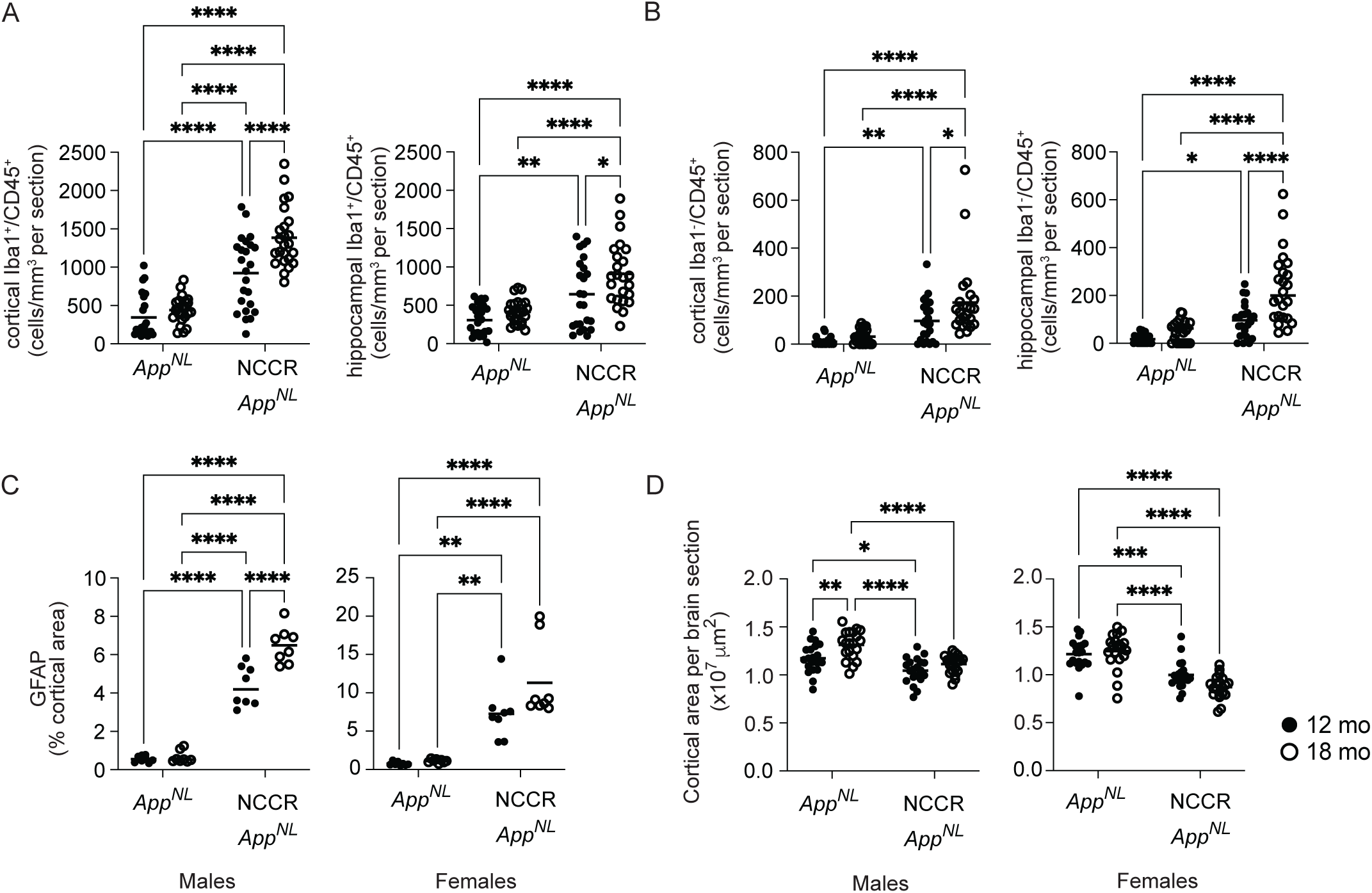
NCCR induces neuroinflammation and cortical atrophy in 12- and 18-month old *App^NL^* mice. (A) Activated microglia was identified with dual labeling for Iba-1 and CD45 in coronal sections from 12- and 18-month old mice. Density of activated microglia was upregulated with NCCR in both cortical and hippocampal areas of the coronal section (two-way ANOVA, genotype effect, p<0.0001). Tukey’s post-hoc analysis showed age-dependent increase in NCCR-*App^NL^* brain samples for both the cortical and hippocampal areas. (B) Brain infiltrating leukocytes were identified as CD45 labeled cells not co-labeled with Iba-1. Density of brain infiltrating leukocytes was increased with NCCR in both the cortical and hippocampal areas (two-way ANOVA, genotype effect, p<0.001). Tukey’s post-hoc analysis showed age-dependent increase in brain leukocyte infiltration induced by NCCR. (C) Astrocytosis was quantified by measuring the cortical area covered by GFAP labeling. Males and females were separately analyzed since there was a significant sex effect (two-way ANOVA, suppl. table 1). Both the males and females showed significant increase in the cortical GFAP labeling with NCCR (two-way ANOVA, genotype effect, p<0.0001; suppl. table 1). Tukey’s post-hoc analysis showed age-dependent increase with NCCR that was statistically significant only in the males (p<0.0001). (D) The effect of NCCR on cortical atrophy was evaluated by measuring the cortical areas in coronal sections. Data from males and females were analyzed separately since the cortical areas for NCCR-*App^NL^* animals showed a sex effect. Both the males and female brain sections showed smaller cortical area with NCCR (two-way ANOVA, genotype effect, p<0.0001). Unlike neuroinflammation, cortical atrophy did not show age-dependent change with NCCR (Tukey’s post-hoc). Complete two-way ANOVA results are provided in suppl. table 2. P-values in the graphs: * p<0.05, ** p<0.01, *** p<0.001, **** p<0.0001. The following numbers of animals were examined: at 12 mo: NCCR-*App^NL^* (n=4M,4F); *App^NL^* (n=4M,4F); at 18 mo: NCCR-*App^NL^* (n=4M,4F); *App^NL^* (n=4M,4F). The numbers of matched coronal sections measured across the animals for the following analyses: Iba1/CD45, n=3; GFAP, n=2; cortical atrophy, n=5. Each data point represents a measure from each section.

With respect to a potential role for NCCR in neurodegenerative processes, both male and female NCCR-*App^NL^* animals displayed significantly smaller cortical areas compared to the *App^NL^* animals at both 12 and 18 months of age (Figure 2D). Coronal brain sections were co-labeled using NeuN and cortical area was evaluated. Data from males and females were analyzed separately since there was a sex difference (Supplementary Figure 3D; Supplementary Table 1). In AD, neurodegeneration and cortical thinning are most strongly correlated with the emergence of tau pathology (Bejanin et al., 2017; Spillantini and Goedert, 2013). Although we did not observe PHF-1+ phospho-tau inclusions in NCCR-*App^NL^* mice, the smaller cortical areas in NCCR-*App^NL^* animals suggest that neuronal cell cycle re-entry induces neurodegeneration in *App^NL^* animals, and the presence of intraneuronal Aβ42 and neuroinflammation in NCCR- *App^NL^* mice could contribute to neurodegeneration. By contrast, we did not observe any statistically significant difference in the cortical areas between 12- and 18-month-old NCCR-*App^NL^* animals in either the males or females, suggesting a lack of progressive cortical atrophy between these time points (Figure 2D). Statistical results for two-way ANOVA analysis by genotype and age related to the main figures are provided in Supplementary Table 2.

### Differential expression analysis highlights upregulation of inflammatory pathways and downregulation of neuronal pathways in NCCR-App^NL^ mice

We leveraged the NanoString Mouse AD gene expression panel to identify a total of 225 differentially expressed genes (DEGs) (adj. p < 0.05) in NCCR-*App^NL^* mice compared to *App^NL^* mice (males and females combined), out of which 153 genes were upregulated (log fold change > 0) and 72 genes were downregulated (log fold change < 0). Microglia-related genes, including *Tyrobp*, *Ctss*, *Cd74*, and complement components *C1qa*, *C1qb*, *C1qc* were highly upregulated in NCCR-*App^NL^* mice compared to *App^NL^* mice, supporting the neuroinflammation-related neuropathological changes observed in these mice. To elucidate the role of these differentially expressed genes, functional enrichment analyses was performed. We identified that *leukocyte transendothelial migration*, *complement and coagulation cascade*, *lysosome*, *platelet activation* and other neuroinflammation-associated pathways were enriched in the list of differentially upregulated genes (Figure 3A), while neuronal related pathways such as *calcium signaling pathway, GABAergic synapse, cholinergic synapse, glutamatergic synapse* were enriched in the differentially downregulated genes (Figure 3B). This suggests that induction of NCCR enhances inflammatory pathways (NES > 2) and suppresses synaptic signaling (NES < -2) (Figure 3C).

**Figure 3.**
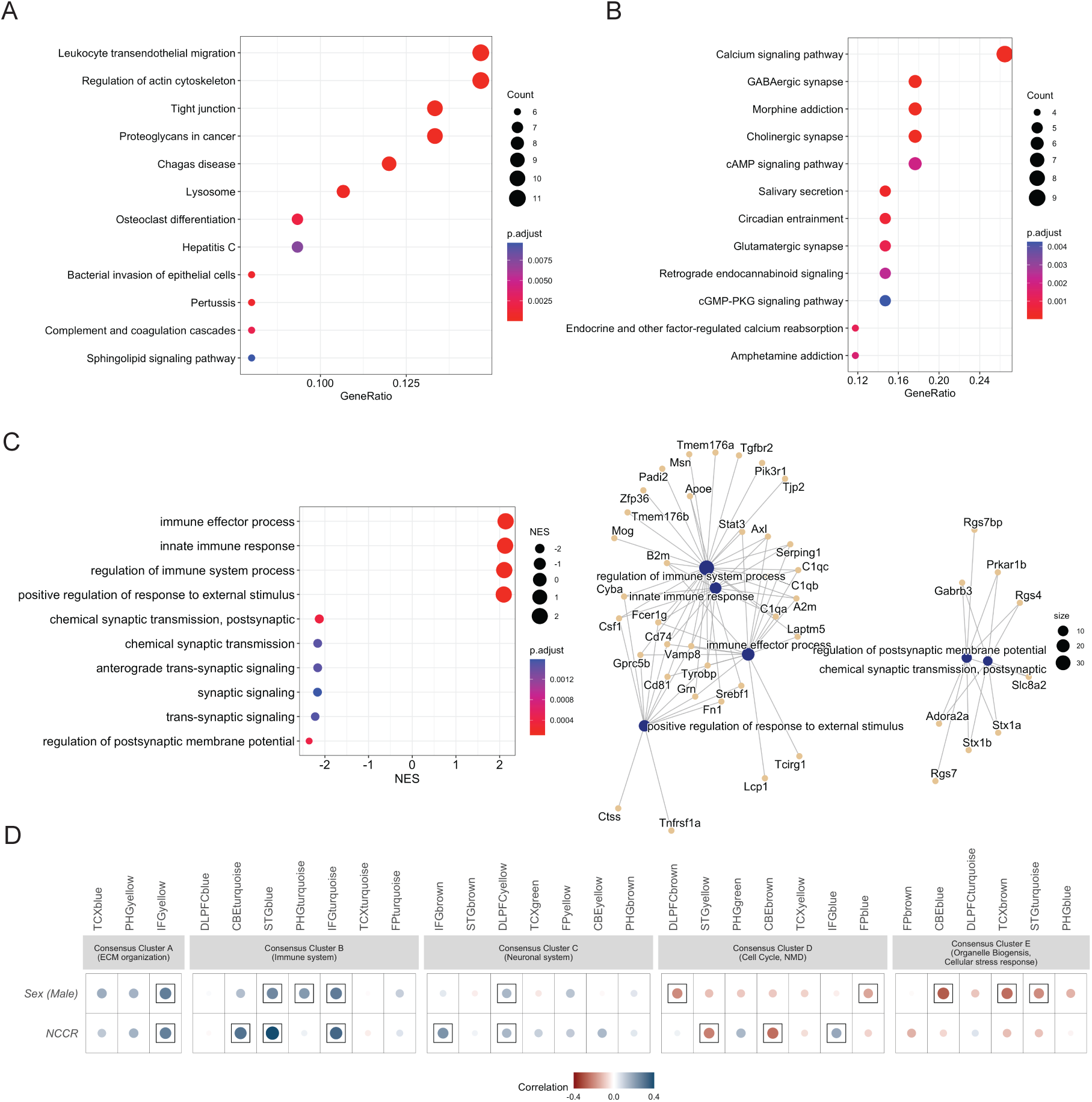
NCCR induces gene expression changes correlated with human AD co-expression modules. Gene expression changes in NCCR-AppNL mice were compared to AppNL mice using NanoString Mouse AD panel to assess the LOAD-relevance of NCCR induced gene expression changes. (A) NanoString gene expression analysis shows significant correlation with various consensus clusters derived from human AD co-expression modules, including that of extracellular matrix. (B,C) Gene set enrichment analysis reveals many functional pathways that are modulated by NCCR for (B) upregulated and (C) down-regulated genes.

To assess the implications of these findings further, we correlated expression changes of NanoString panel genes observed in NCCR-*App^NL^* mice compared to *App^NL^* mice with expression changes of these genes in 5xFAD mice compared to wild type B6 mice (Preuss et al., 2020). There was a significant positive correlation (R=0.26, p < 1E-13) for the extent and directionality of gene expression changes between NCCR-induced changes and 5xFAD mice (Supplementary Figure 4A). However, there were distinct differences in the expression pattern between the two mouse models. Specifically, we noticed upregulation of above-mentioned microglia and complement components genes in both NCCR and 5xFAD mice, yet the changes in expression were higher in NCCR mice as compared to 5xFAD mice, suggesting NCCR induces greater inflammatory responses despite the absence of Aβ plaques (Supplementary Figure 4A). We also found that multiple AD risk genes such as *CD74*, *TYROBP*, *CSF1* and *CTSS* were upregulated in both NCCR and 5XFAD mice and in immune system-related human co-expression modules (Supplementary Figure 4B).

In contrast, many of the genes significantly altered by NCCR were unaffected in 5XFAD mice, including *Cd74, Ctss* and *C1qb* genes. On the other hand, *Trem2* gene expression was not significantly upregulated (logFC=0.13; FDR > 0.05) with NCCR, whereas it was significantly upregulated in the 5XFAD mice compared to B6 (logFC=0.40; FDR < 0.05).

### NCCR in App^NL^ mice correlates with distinct human co-expression modules in brain region and pathway-specific manner

To assess the relevance of NCCR in LOAD, we performed a correlation analysis between human post-mortem co-expression modules (Wan et al., 2020) and mouse brain transcriptomic data from the NanoString Mouse AD panel (Preuss et al., 2020). The effects of NCCR on the brain transcriptome of *App^NL^* mice showed significant positive correlations (Pearson’s correlation coefficient > 0.25; p<0.05) with human co-expression modules (IFGyellow) in Consensus Cluster A, which is enriched for genes associated with extracellular matrix (ECM) in the inferior frontal gyrus (IFG) brain region (Figure 3D). Furthermore, NCCR displayed significant positive correlations (Pearson’s correlation coefficient > 0.3; p<0.05) with immune-related human co-expression modules (CBEturquoise, STGblue, and IFGturquoise) in the IFG, superior temporal gyrus (STG), and cerebellum in Consensus Cluster B (Figure 3D). NCCR also showed significant positive correlations (Pearson’s correlation coefficient > 0.2; p<0.05) with neuronal co-expression modules enriched for synaptic signaling (IFGbrown and DLPFCyellow) in Consensus Cluster C (Figure 3D). Interestingly, NCCR displayed a significant positive correlation (Pearson’s correlation coefficient = 0.2; p<0.05) with human module (IFGblue) in the IFG as well as significant negative correlations (Pearson’s correlation coefficient < -0.2; p<0.05) with human modules (STGyellow and CBEbrown) in the STG and cerebellum in Consensus Cluster D that is enriched for transcripts associated with cell cycle, myelination, and glial development (Figure 3D). By contrast, NCCR did not show a significant positive correlation (p > 0.05) with any modules in Consensus Cluster E, which is enriched for transcripts associated with organelle biogenesis and cellular stress response pathways in multiple brain regions. Overall, we observed overlaps with multiple human co-expression modules associated with distinct disease processes and molecular pathways, suggesting the potential translational relevance of NCCR-induced pathogenesis for modeling LOAD.

To further dissect the pathways driving these module-level results, we extracted genes exhibiting directional coherence between the effects of NCCR and changes in expression in AMP-AD modules, followed by KEGG and REACTOME pathway enrichment analysis to elucidate the role of these disease-related genes. A total of 128 genes from the NCCR-mouse Nanostring datset, such as *Aldh2, Aldh6a1, Arhgap5, Bcl2, Cldn10, Itgb5, Lamb2, Rab31*, and *Sox9*, that exhibited directional coherence with gene expression changes in ECM associated human modules in Consensus Cluster A were identified (Supplementary Figure 5A). Many of these genes were upregulated for the NCCR effect, and the KEGG pathway analysis identified enrichment of *regulation of actin cytoskeleton*, *fatty acid degradation*, *ECM-receptor interaction*, *focal adhesion*, and *leukocyte transendothelial migration* associated pathways in these coherent gene sets (Supplementary Figure 5B). Next, we identified a total of 201 genes altered by NCCR exhibiting directional coherence with immune associated genes from human modules in Consensus Cluster B, including microglia related genes *Trem2, Tyrobp*, and complement components *C1qa, C1qb, C1qc* (Supplementary Figure 6A). These genes were upregulated in both human modules and NCCR mice, and enriched for *leukocyte transendothelial migration*, *osteoclast differentiation*, *chagas disease*, and *tight Junction* associated pathways (Supplementary Figure 6B). Similarly, we identified a total of 222 genes from NCCR effect exhibiting directional coherence with neuronal associated genes from human co-expression modules in Consensus Cluster C, such as *Slc6a17, Gabarapl1, Calm3, Mapt*, and *Ppp3cb* (Supplementary Figure 7A). These disease-related genes were downregulated in both NCCR mice and human modules in Consensus Cluster C, and were enriched for *Alzheimer’s disease*, *calcium signaling pathway*, *GABAergic synapse* and neurodegeneration associated pathways (Supplementary Figure 7B).

Although NCCR showed a significant positive correlation with only one cell cycle-associated module (IFGblue) in Consensus Cluster D, a total of 247 genes altered by NCCR exhibited directional coherence with cell cycle and myelination associated genes in human AD modules in Consensus Cluster D such as *Bin1, Abca2, Picalm* and *Cdk18* (Supplementary Figure 8A). These disease-related genes were enriched for *PI3K-Akt signaling, RHO GTPase cycle and CDC42 GTPase cycle* associated pathways (Supplementary Figure 8B-C, Supplementary File X). Finally, we identified a total of 141 NCCR-related genes exhibiting directional coherence with genes from human co-expression modules in Consensus Cluster E such as *Dnajb6, Hsp90aa1, Hsph1, Hsbp1* and *Psma5 (Supplementary Figure 9A)*. These genes were enriched for *heat-shock response*, *regulation of mitotic cell cycle*, *protein processing in endoplasmic reticulum*, and *proteasome* associated pathways (Supplementary Figure 9B).

### NCCR mice show similarities and differences with existing FAD and LOAD mouse models

Next, we systematically compared the NCCR mice with previously studied 5XFAD and late-onset risk variant mice at the pathway level (Preuss et al., 2020). NanoString AD panel genes were ranked based on regression coefficients calculated for NCCR, 5XFAD, APOE4 and Trem2.R47H mice, and gene set enrichment analysis was performed using both KEGG and REACTOME pathway libraries (Subramanian et al., 2005). We identified multiple similar REACTOME pathways in both NCCR and 5XFAD mice. Pathways including *fatty acid metabolism*, *extracellular matrix organization*, and *immune system* were upregulated (NES > 0) (Figure 4A), whereas neuronal system-associated pathways were downregulated (NES < 0) (Figure 4A). However, we identified some divergent REACTOME pathways such as *cell cycle/mitotic*, *MAPK1/MAPK3 signaling*, *cell cycle checkpoints* and *signaling by interleukins*, which were downregulated (NES < 0) in NCCR-AD mice but upregulated (NES > 0) in 5XFAD and other LOAD models (Figure 4A). KEGG pathways also showed many similarities between NCCR mice and other commonly used AD mouse models (Figure 4B). For example, *Lysosome* pathway is upregulated while second messenger signaling pathways such as *Calcium signaling, cAMP signaling,* and *Oxytocin signaling* pathways are down regulated in the four different mouse models compared (Figure 4B). Therefore, our findings suggest that NCCR can impart many pathway changes that are either similar to or distinct from other EOAD and LOAD mouse models.

**Figure 4.**
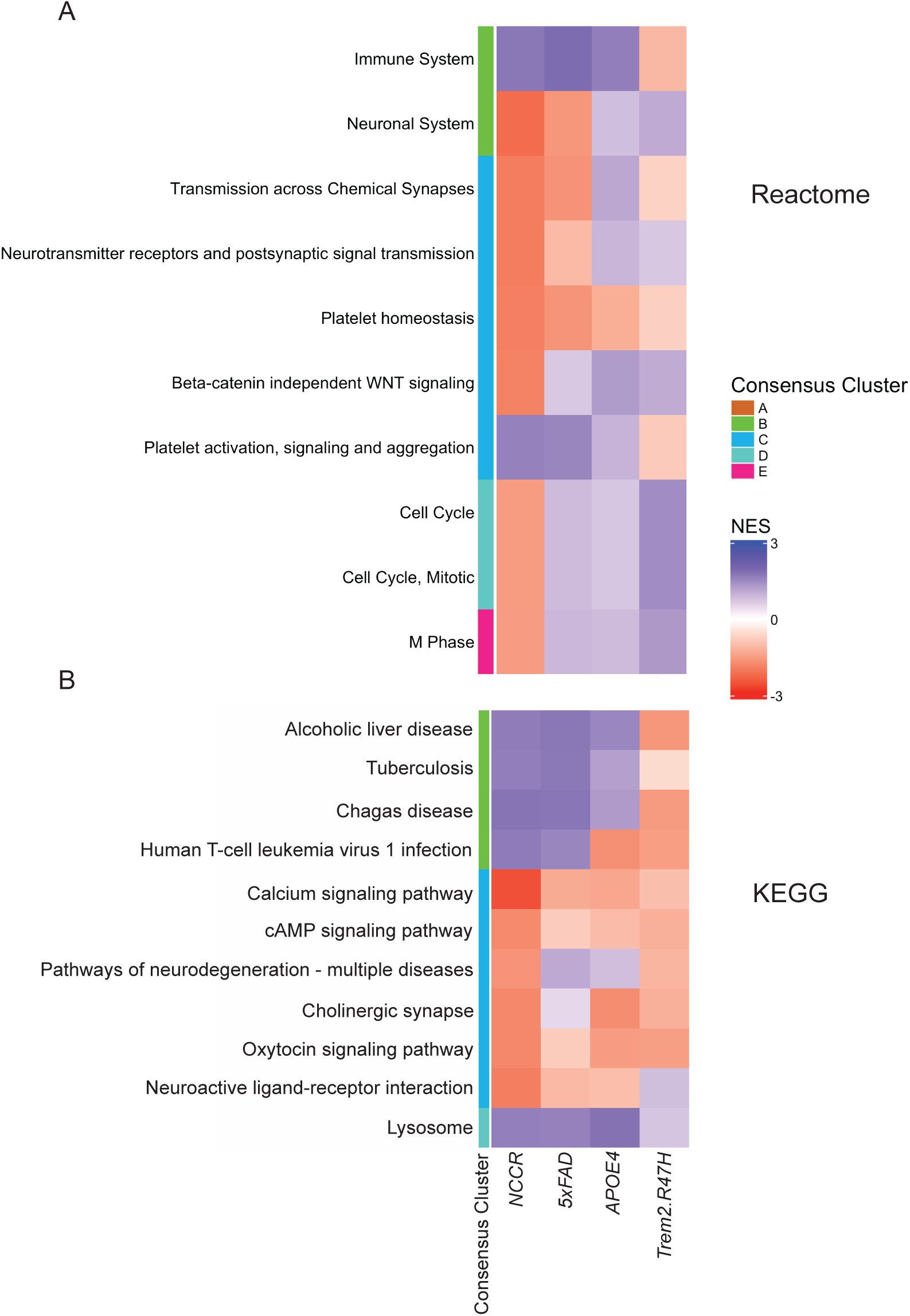
Pathway comparison of NCCR mice with other AD mouse models based on NanoString AD panel genes. Heat map for top gene set enriched pathways identified from KEGG and REACTOME libraries is shown. Normalized enrichment scores (NES) for ranked genes demonstrate similar and divergent pathways between NCCR mice and other AD mouse models such as 5xFAD, APOE4, and TREM2 mice.

### Suppressing NCCR at a middle age reduces AD-associated neuropathological features at a later age

A doxycycline (dox) diet can be used to selectively manipulate neuronal cell cycle re-entry through the conditional expression of SV40T in our unique NCCR mouse model (Park and Barrett, 2020). In order to directly evaluate the role of NCCR in the progression of AD-associated neuropathological features, we put the NCCR-*App^NL^* and NCCR-*App^NL-F^* animals back on continuous dox diet again starting at 12 months of age and their AD-associated neuropathological features were evaluated at 18 months of age (i.e., 11 months of regular diet followed by 6 months of dox diet). We hypothesized that if disease progression is NCCR-dependent, halting NCCR should prevent disease progression. This is also an important question regarding the therapeutic viability of manipulating cell cycle pathways as a disease-modifying strategy.

As anticipated, 18-month old NCCR-*App^NL^* animals that are put back on dox diet did not show any SV40T or PCNA demonstrating successful suppression of NCCR (Supplementary Figure 2B). Additionally, we included the NCCR-*App^NL-F^* animals for evaluating the effect on PHF-1+ phospho-tau lesions since these features were not observed in the NCCR-*App^NL^* animals. Evaluation of the 18-month old animals showed that all of the AD-associated neuropathological features are diminished in the NCCR-*App^NL^* and NCCR-*App^NL-F^* mice that are put back on continuous dox diet starting at 12 months of age (Figure 5; accompanying two-way ANOVA results are provided in Supplementary Table 3). For instance, the number of cells displaying intraneuronal Aβ(x-42) accumulation was reduced by 72 percent in the 18-month old NCCR-*App^NL^* animals that were put back on dox diet compared to the 18-month old NCCR-*App^NL^* animals that were continuously maintained on regular diet (Figure 5A). The number of intraneuronal Aβ(x-42)-immunopositive neurons were also 50 percent lower compared to 12-month old NCCR-*App^NL^* animals maintained on a regular diet (Figure 5A). These findings indicate that the progressive increase in intraneuronal Aβ(x-42) observed in this model is NCCR-dependent.

**Figure 5.**
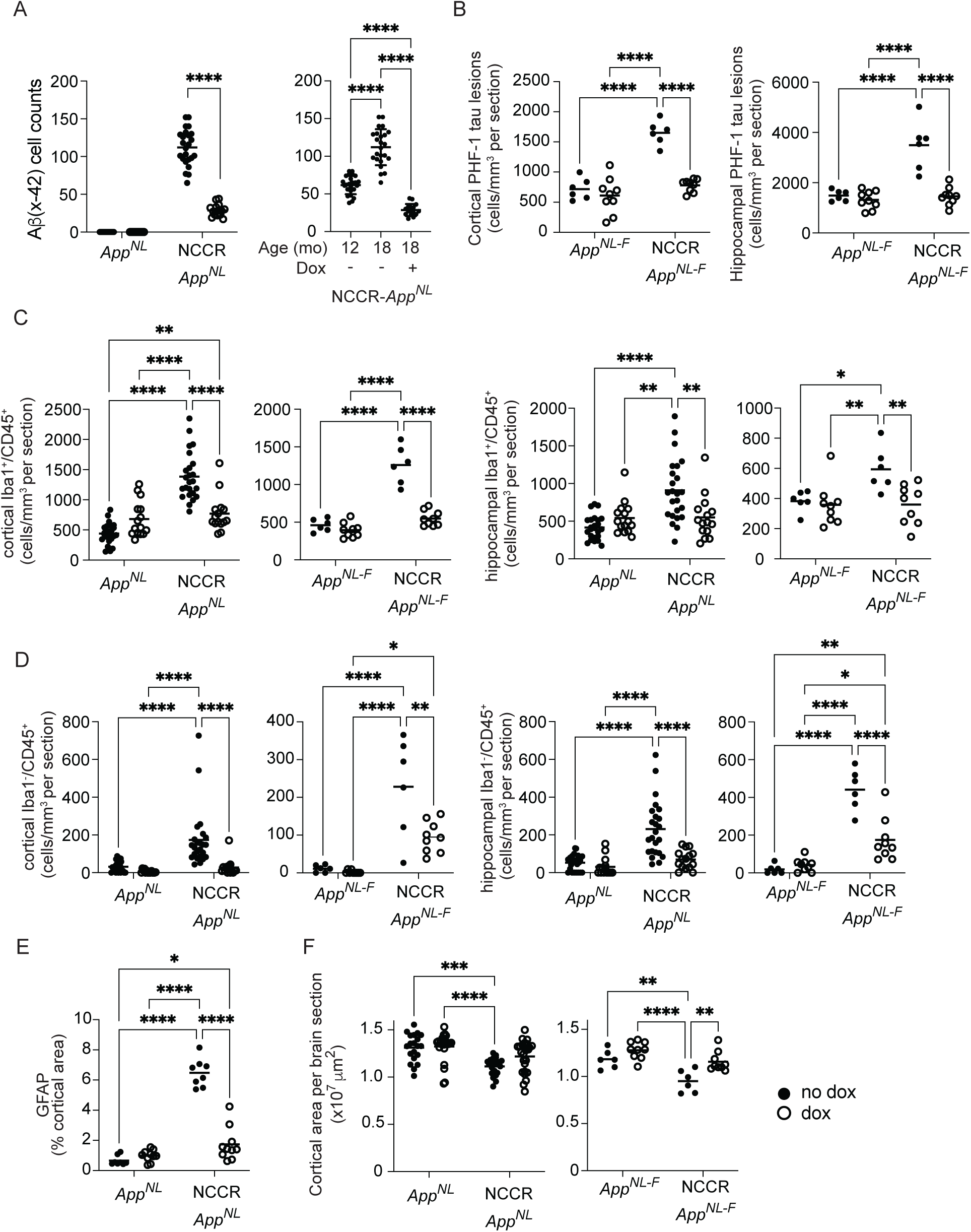
NCCR drives the AD-associated neuropathological features. The effect of suppressing the NCCR on AD pathologies was evaluated at 18 months of age after putting the animals back on continuous dox diet starting at 12 months of age. NCCR-*App^NL^* mice were evaluated for intraneuronal Aβ(x-42) and NCCR-*App^NL-F^* mice were evaluated for PHF-1 phospho-tau lesions. The NCCR animals that were put back on dox diet showed lower number of (A) cells with intraneuronal Aβ (x-42), (B) cells bearing PHF-1 phospho-tau lesions, (C) activated microglia, (D) brain infiltrating leukocytes. (E) Cortical area was greater in both NCCR-*App^NL^* and NCCR-*App^NL-F^* animals that were put back on dox diet compared to 18-month old animals that were continuously maintained on a regular diet. However, the increase was statistically significant only in NCCR-*App^NL-F^* animals that were put back on dox diet (Tukey’s post-hoc test, p<0.01). Furthermore, both NCCR-*App^NL^* and NCCR-*App^NL-F^* animals that were put back on dox diet showed similar cortical areas as the respective *App^N-L^* and *App^NL-F^* animals (Tukey’s post-hoc, p>0.05). (F) Astroyctosis was also lower in the NCCR-*App^NL^* animals that were put back on dox diet (Tukey’s post-hoc, p<0.0001). Complete two-way ANOVA results are provided in suppl. table 2. P-values in the graphs: * p<0.05, ** p<0.01, *** p<0.001, **** p<0.0001. The following numbers of 18-month old NCCR-*App^NL^* and *App^NL^* animals were examined: put back on dox, NCCR-*App^NL^* (n=5M), *App^NL^* (n=5M); regular diet (no dox), NCCR-*App^NL^* (n=4M), *App^NL^* (n=4M). The numbers of matched coronal sections were measured across the animals for the following analyses: Aβ(x-42), n=3; Iba1/CD45, n=3; GFAP, n=2; cortical atrophy, n=5. The following numbers of 18-month old NCCR-*App^NL-F^* and *App^NL-F^* animals were examined: put back on dox, NCCR-*App^NL-F^* (n=3F), *App^NL-F^* (n=2M, 1F); regular diet (no dox), NCCR-*App^NL-F^* (n=1M, 1F), *App^NL-F^* (n=1M, 1F). The numbers of matched coronal sections measured across the animals for the following analyses: Iba1/CD45, n=3; PHF-1, n=3; cortical atrophy, n=3. Each data point represents a measure from each section.

Similarly, there was a 50-55% reduction in the number of cortical neurons bearing PHF-1+ phospho-tau epitopes in the cortex and hippocampus of 18-month old NCCR-*App^NL-F^* animals that were put back on dox diet starting at 12 months of age compared to the 18-month old NCCR-*App^NL-F^* animals that were continuously maintained on regular diet (Figure 5B). Furthermore, post hoc analysis showed that the number of PHF-1+ phospho-tau lesions in the 18-month old NCCR-*App^NL-F^* mice that were put back on dox diet were not different from 18-month old *App^NL-F^* animals that were either continuously maintained on regular diet or put back on dox diet starting at 12 months of age (Figure 5B). Therefore, halting NCCR inhibited the accrual of tau pathology in this model.

Microglia activation and brain leukocyte infiltration were also suppressed in the cortex and hippocampus of both NCCR-*App^NL^* and NCCR-*App^NL-F^* mice at 18-months of age after being put back on dox diet in comparison to the age-matched animals that were continuously maintained on regular diet (Figure 5C and D). These levels were not statistically different from those observed in age-matched *App^NL^* or *App^NL-F^* animals that were also maintained on dox diet starting at 12 months of age (Figure 5C and D). Additionally, cortical GFAP immunoreactivity was decreased by 76% in 18-month old NCCR-*App^NL^* mice on the dox diet compared to 18-month old NCCR-*App^NL^* animals maintained on a regular diet (Figure 5E). Furthermore, measures of these neuroinflammatory features in NCCR animals put back on dox diet were not different from those measured in respective *App^NL^* and *App^NL-F^* control groups (Figure 5C,D and E). Thus, halting NCCR helped normalize neuroinflammatory changes in this model.

Finally, inhibiting NCCR via the dox diet also helped mitigate cortical atrophy in the NCCR-*App^NL-F^* and NCCR-*App^NL^* animals. The cortical areas for the 18-month old NCCR-*App^NL^* animals put back on dox diet were greater than age-matched NCCR-*App^NL^* animals continuously maintained on regular diet, but this difference was not statistically different (Figure 5F). However, the cortical areas for the 18-month old *App^NL^* mice were greater when compared to the 18-month old NCCR-*App^NL^* animals maintained on regular diet, but not compared to the 18-month old NCCR-*App^NL^* animals that were put back on dox diet (Figure 5F). Similarly, the cortical areas for the 18-month old *App^NL-F^* mice were greater when compared to the 18-month old NCCR-*App^NL-F^* animals maintained on regular diet, but not compared to the 18-month old NCCR-*App^NL-F^* animals that were put back on dox diet (Figure 5F). Additionally, 18-month old NCCR-*App^NL-F^* animals that were put back on dox diet showed larger cortical areas compared to 18-month old NCCR-*App^NL-F^* that were continuously maintained on regular diet (Figure 5F). Hence, halting NCCR helped prevent cortical atrophy in this model. These findings also support our previous findings that showed age-dependent cortical atrophy in the NCCR-*App^NL-F^* animals (Barrett et al., 2021), and these observations highlight the additional impact of tau pathology on neurodegeneration.

### LCM-microarray analysis reveals AD-associated pathway changes in SV40T-positive neurons

Inhibiting NCCR via a back-on-dox diet manipulation showed that neuronal SV40T expression drives AD-like disease progression in a cell-autonomous manner. SV40T is expressed in approximately 5% of the cortical neurons in our NCCR mouse model (Barrett et al., 2021). In this regard, laser capture microdissection (LCM)-microarray analysis of SV40T-immunopositive versus naive neurons (SV40T-negative) from 4-month-old animals showed 73 DEGs (fold-change ≥ |2|, adj. p < 0.05), of which 47 were upregulated and 26 were downregulated. KEGG and REACTOME pathway analyses showed that the upregulated genes were enriched for pathways associated with *autophagy* (KEGG, adj. p = 1.45E-05; REACTOME, adj. p = 5.74E-05), and *detoxification of ROS pathway* (REACTOME, adj. p = 0.02) (Supplementary Figure 10A). The downregulated genes were enriched for pathways associated with *Glycolysis and gluconeogenesis* (KEGG, adj. p = 0.0038; REACTOME, adj. p = 0.0082), *HIF-1 signaling* (KEGG, adj. p = 0.0096), *glutamatergic synapse* (KEGG, adj. p = 7.65e-05), *Class C/3(metabotropic glutamate /pheromone receptors*) pathway, REACTOME, adj. p = 0.0014), and *neurexin/neuroligin* (REACTOME, adj. p = 0.017) (Supplementary Figure 10B).

There were 8 genes on LCM-microarray that overlapped with NanoString genes. Of these 8, only A2M, GLUD1, and RGS7 were differentially regulated on NanoString gene panel. A2M was upregulated in both LCM and NanoString analysis, whereas GLUD1 and RGS7 were inversely related. GLUD1 was downregulated in LCM-microarray but upregulated in NanoString, whereas RGS7 was upregulated in LCM-microarray but downregulated in NanoString. Additionally, LCM-microarray showed increases in MLH1 (DNA mismatch repair, 4.9-fold), ERCC1 (DNA excision repair, 2.8-fold), 1N4R tau (2.8-fold) transcripts in SV40T+ neurons that were notable, especially since 1N4R tau isoform transcript is the most abundant of the six tau isoforms found in human brain.

## Discussion

We have previously demonstrated that NCCR-AD mice display many LOAD-like neuropathological features (Barrett et al., 2021; Park et al., 2007; Park and Barrett, 2020). Here, we extended these observations by employing an integrated and complementary transcriptomic, bioinformatic, and pathological analytical approach to demonstrate the translational relevance of NCCR-AD mouse models for identifying mechanisms and potential preclinical therapeutic targets for LOAD. Furthermore, by putting the animals back on dox diet starting at 12 months of age, we confirmed that NCCR drives the age-related neuropathological changes observed in the 18-month old NCCR-*App^NL^* and NCCR-*App^NL-F^* mice, supporting the concept that targeting NCCR can modify AD progression.

Our previous work showed that when combined with our NCCR mouse model, many of the AD-related neuropathological features are enhanced in *App^NL-F^* KI mice (Barrett et al., 2021). However, the effect of NCCR on Aβ processing was unclear in that study likely due to excessive levels of Aβ_42_ produced in *App^NL-F^* KI animals. By crossing our NCCR mice with *App^NL^* KI mice, we clearly demonstrate that NCCR promotes the formation of intraneuronal Aβ(x-42). Furthermore, NCCR induces neuroinflammation, brain leukocyte infiltration, and neurodegeneration in *App^NL^* animals that otherwise do not display neuropathological changes. Notably, despite the age-dependent increase in intraneuronal Aβ(x-42), NCCR-*App^NL^* mice did not display Aβ plaques or PHF-1+ phospho-tau lesions at either 12 or 18 months of age, in contrast to our previous study showing increased PHF-1+ phospho-tau lesions with NCCR in *App^NL-F^* mice (Barrett et al., 2021).

The lack of Aβ plaques and PHF-1+ phospho-tau lesions in these mice are likely due to high levels of Aβ_40_ in the *App^NL^* mice. Aβ_42_ peptides have been found to be more neurotoxic (Klein et al., 1999) and more prone to aggregation than Aβ_40_ due to their ability to self-assemble into oligomers and fibrils, and as result contribute significantly to amyloid plaque formation (Kayed et al., 2003; Marshall et al., 2016). Additionally, it has been shown that Aβ_40_ may inhibit plaque formation (Kim et al., 2007) and that Aβ_40_/ Aβ_42_ ratios modulate the extent of tau pathology (Kwak et al., 2020).

Although much knowledge regarding the role of Aβ in AD has been gained using mutant *App* FAD mouse models, the lackluster results from Aβ-targeted clinical trials highlight the need for alternative pathogenic mechanisms of AD. Our neuropathological and gene expression profiling data provide compelling new data for the translational relevance of NCCR mouse model (NCCR-AD) for LOAD. NCCR-AD mice display gene expression changes that display significant concordance with functional pathways associated with AD and, as such, they hold great promise for revealing novel preclinical clues to disease pathophysiology. For example, NCCR sufficiently drives AD-relevant neuroinflammation processes, as alpha2-macroglobulin (A2M) expression is upregulated in both the SV40T-expressing neurons (LCM-microarray data) as well as globally in the whole brains of late-stage NCCR animals (NanoString mouse AD panel) in the absence of Aβ plaques. A2M, a pan-protease inhibitor, is expressed in neurons in AD brains but not in normal aged brains (Bauer et al., 1991). Furthermore, expression of A2M has been identified as a potential blood biomarker for neurodegeneration and inflammation (Varma et al., 2017) and positively correlated with AD disease progression (Varma et al., 2017). Additionally, our previous work has shown that gliosis is one of the earliest changes induced by NCCR (Park and Barrett, 2020). Therefore, increased A2M expression in SV40T-postive neurons could be associated with neuroinflammation in the NCCR-AD mice (Naseraldeen et al., 2021). Additionally, intraneuronal Aβ42 also has been shown to be neuroinflammatory (Welikovitch et al., 2020). Taken together, these data show that NCCR can induce neuronal changes that can elicit neuroinflammatory responses, although the exact processes remain to be elucidated.

Another interesting result from our NanoString nCounter mouse AD panel analysis was the significant positive correlation between NCCR mice and human co-expression modules associated with extracellular matrix (ECM) organization in LOAD processes. On the other hand, 5xFAD mice which are the most commonly used AD mouse model do not show LOAD-relevance for this co-expression module (Preuss et al., 2020). The NanoString data suggest that aberrant NCCR can trigger AD-associated changes in the ECM that is not present in 5xFAD mice. Since ECM plays an important role in the inflammation process by modulating immune cell response and behavior (Sorokin, 2010), the increase in neuroinflammation and brain leukocyte infiltration could arise from ECM changes in NCCR mice. Also, it has been suggested that ECM changes are present in early stages of AD, and these changes could contribute to vascular changes (Davis and Senger, 2005; Lepelletier et al., 2017). Hence, an evaluation of vascular function in the NCCR-AD mice is warranted to determine whether these mice also mirror vascular dysfunction associated with AD..

One of the most interesting neuropathological features in the NCCR-AD mice is the presence of PHF-1+ tau lesions in the absence of human *MAPT* or tau mutations. However, unlike the NCCR-*App^NL-F^* mice, NCCR-*App^NL^* mice did not display Aβ plaques or PHF-1+ tau lesions. Our data lend support to the notion that the presence of Aβ plaques is necessary for PHF-1+ tau lesions. AD is believed to be a secondary tauopathy where Aβ pathology precedes the appearance of tau pathology (Jack et al., 2018). It has been shown that Aβ plaque enhances tau seeding *in vivo* and that the Aβ42/40 ratio drives tau pathology in 3D human neural cell culture (He et al., 2018; Kwak et al., 2020). Furthermore, the relationship between Aβ and tau pathologies is demonstrated by concomitant reduction in CSF p-tau levels when brain Aβ load is depleted using Aβ antibodies in AD patients (Shi et al., 2022). Therefore, NCCR-AD mice distinctly model the relationship between Aβ plaques and AD tau pathology *in vivo*. Additionally, our LCM-microarray analysis demonstrates that 1N4R tau isoform transcript, which is the most abundant of the six tau isoforms found in human brain, is upregulated in SV40T-positive neurons. It has been shown that transgenic overexpression of wildtype 1N4R tau isoform can induce tau pathologies in mice (Wheeler et al., 2015), and It has been suggested that RNA splicing is dysregulated in tauopathies (Apicco et al., 2019). Further evaluation in NCCR-AD mice on the role of neuronal dysregulation in RNA splicing and its impact on the disease could provide insight into AD tau pathogenesis.

Findings from our study demonstrates that NCCR is the primary pathogenic driver of AD-related neuropathological features in our mice. Functional pathway enrichment analysis of neuronal gene expression changes highlighted in our LCM-microarray data shows that numerous functional processes thought to be associated with AD pathogenic processes are altered in 4-month-old NCCR-AD mice, including *glucose metabolism* and *autophagy* pathways. Furthermore, neuronal upregulation of *MLH1* and *ERCC1* likely reflect DNA damage repair responses resulting from aberrant NCCR, which is suggested by the presence of γ-H2AX signal in these mice (Barrett et al., 2021). These changes are associated with downregulation of pre- and post-synaptic genes associated with excitatory neurons, which likely reflects the nature of Camk2a-driven NCCR. It has been shown that changes associated with genomic damage are present in excitatory neurons in AD brains (Miller et al., 2022). Thus, our findings suggest that NCCR-mediated oxidative and genotoxic stress promote AD pathogenesis including Aβ-and tau-pathologies.

## Conclusions

The majority of AD cases are sporadic and the risk of developing the disease is influenced by numerous environmental and genetic factors. However, finding a cure for the disease has been challenging since so many cell types and cellular dysfunctions seemingly contribute to the disease while the pathogenic mechanisms remain unknown. Our NCCR-AD mice represent an AD mouse model in which global metabolic dysfunction in vulnerable neurons can drive many LOAD-relevant changes in gene networks and neuropathological features *in vivo*. In summary, our NCCR-AD mice represent a novel and exciting translationally relevant AD mouse model for identifying new drugs and treatments targeting neuronal dysfunction driving AD pathogenesis.

## Funding

This work was supported by the National Institutes of Health grant AG060144 to K.H.J.P.

## Abbreviations

IFG: inferior frontal gyrus
STG: superior temporal gyrus
CBE: cerebellum
DLPFC: dorsolateral prefrontal cortex
FP: frontal pole
TCX: temporal cortex
PHG: parahippocampal gyrus
NCCR: neuronal cell cycle re-entry
ECM: extracellular matrix
LOAD: late-onset Alzheimer’s disease
FAD: familial Alzheimer’s disease
AD: Alzheimer’s disease
AMP-AD: accelerating medicines partnership program for AD
MODEL-AD: Model Development and Evaluation for LOAD
NES: normalized enrichment score
DEG: differentially expressed genes
APP: amyloid precursor protein
NFT: neurofibrillary tangle
LCM: laser capture microdissection
TRE: tetracycline response element
tTA: tetracycline transactivator
Dox: doxycycline
GSEA: gene set enrichment analysis

## Declarations of interest

none

### Disclosures

The authors report no competing financial or other interests.

### Contributors

K.A.S..: conceptualization, neuropathological data collection and analysis.

R.S.P.: gene expression data analysis, bioinformatics, writing the paper.

A.W.: neuropathological data collection and analysis.

J.S.B.: LCM-microarray data collection and analysis.

S.E.C.: LCM-microarray data collection and analysis, writing the paper.

G.W.C.: gene expression data analysis, bioinformatics, writing the paper.

K.H.J.P.: conceptualization, data analysis, supervision, project administration, writing the paper, and funding acquisition.

## Acknowledgements

We would like to thank Carol Stevens for help with mouse colony management.

## Notes

### Competing Interest Statement

The authors have declared no competing interest.

