## supplemental figures for "Aberrant neuronal cell cycle re-entry induces late-onset Alzheimer’s disease relevant neuropathological and gene expression changes"

Suppl. Figure 1 v1

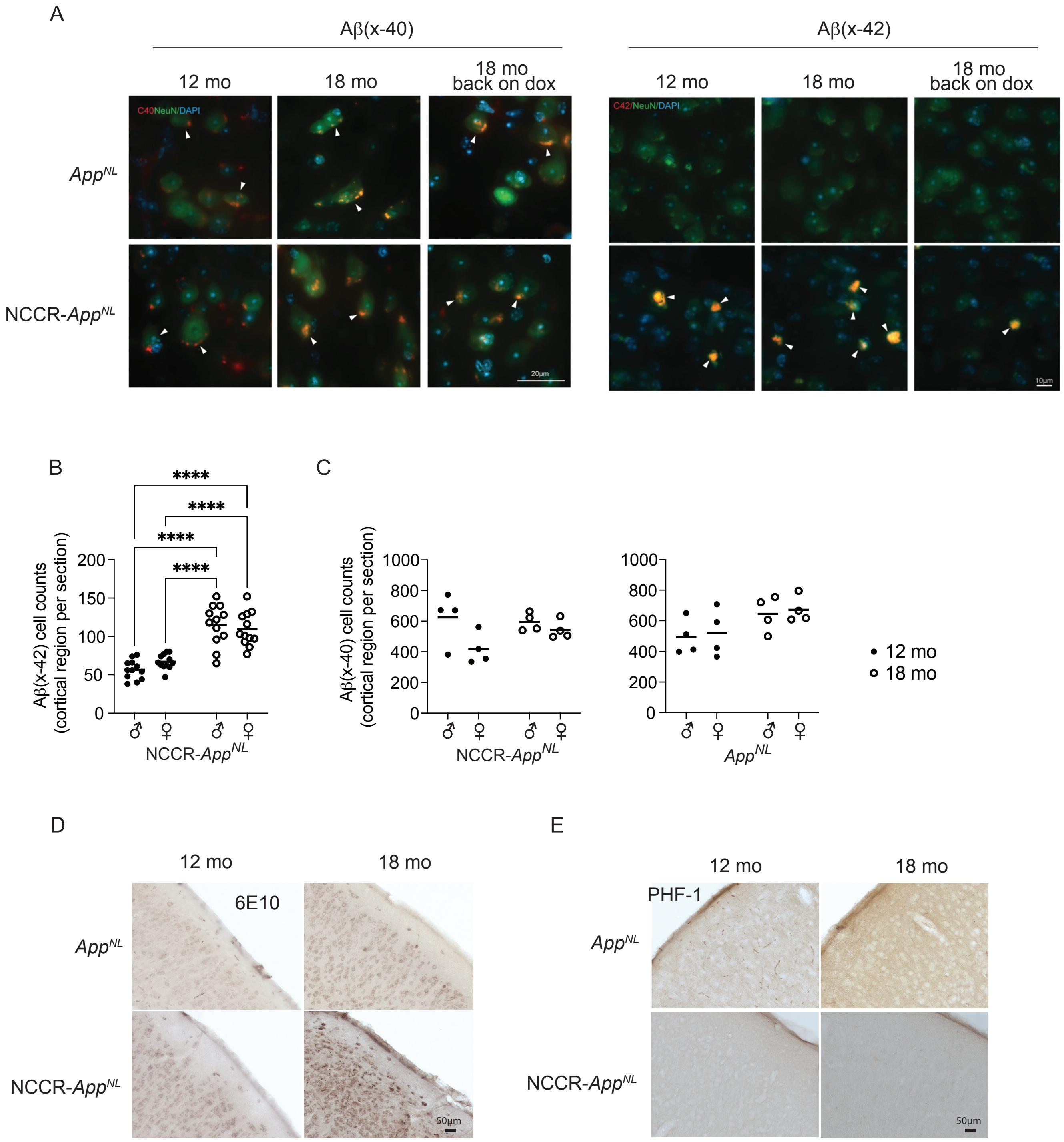

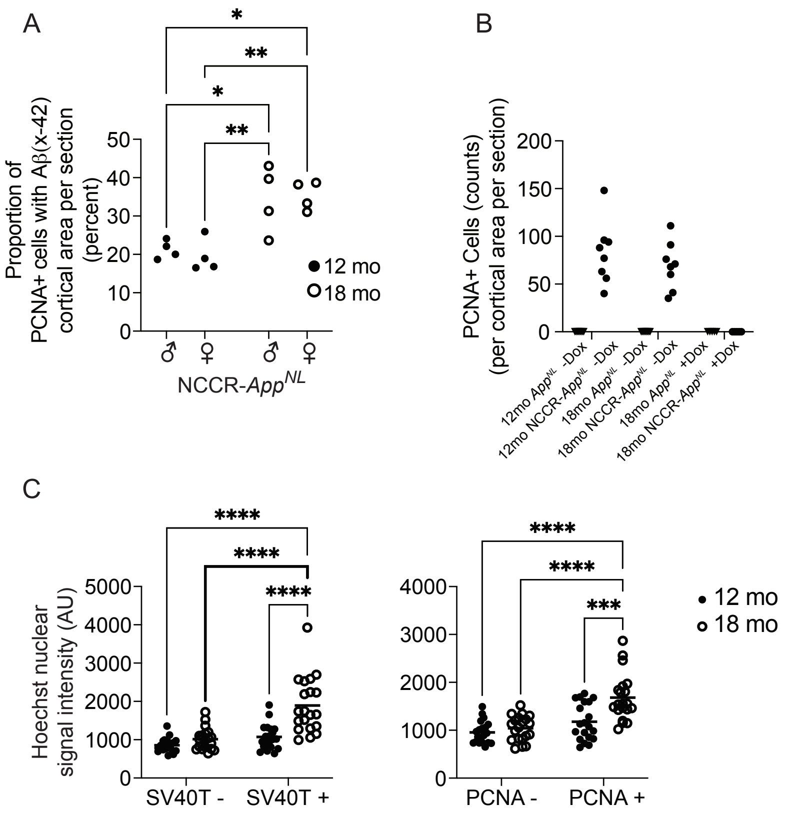
A
*
**
*
**
Proportion of PCNA+ cells with Aβ(x-42) cortical area per section (percent)
50
40
30
20
10
0
12 mo
18 mo
♂ ♀ ♂ ♀
NCCR-AppNL
B
PCNA+ Cells (counts) (per cortical area per section)
200
150
100
50
0
12mo AppNL -Dox
12mo NCCR-AppNL -Dox
18mo AppNL -Dox
18mo NCCR-AppNL -Dox
18mo AppNL +Dox
18mo NCCR-AppNL +Dox
C
****
****
****
Hoechst nuclear signal intensity (AU)
5000
4000
3000
2000
1000
0
SV40T -
SV40T +
****
****
***
4000
3000
2000
1000
0
PCNA -
PCNA +
12 mo
18 mo

Suppl. Figure 3

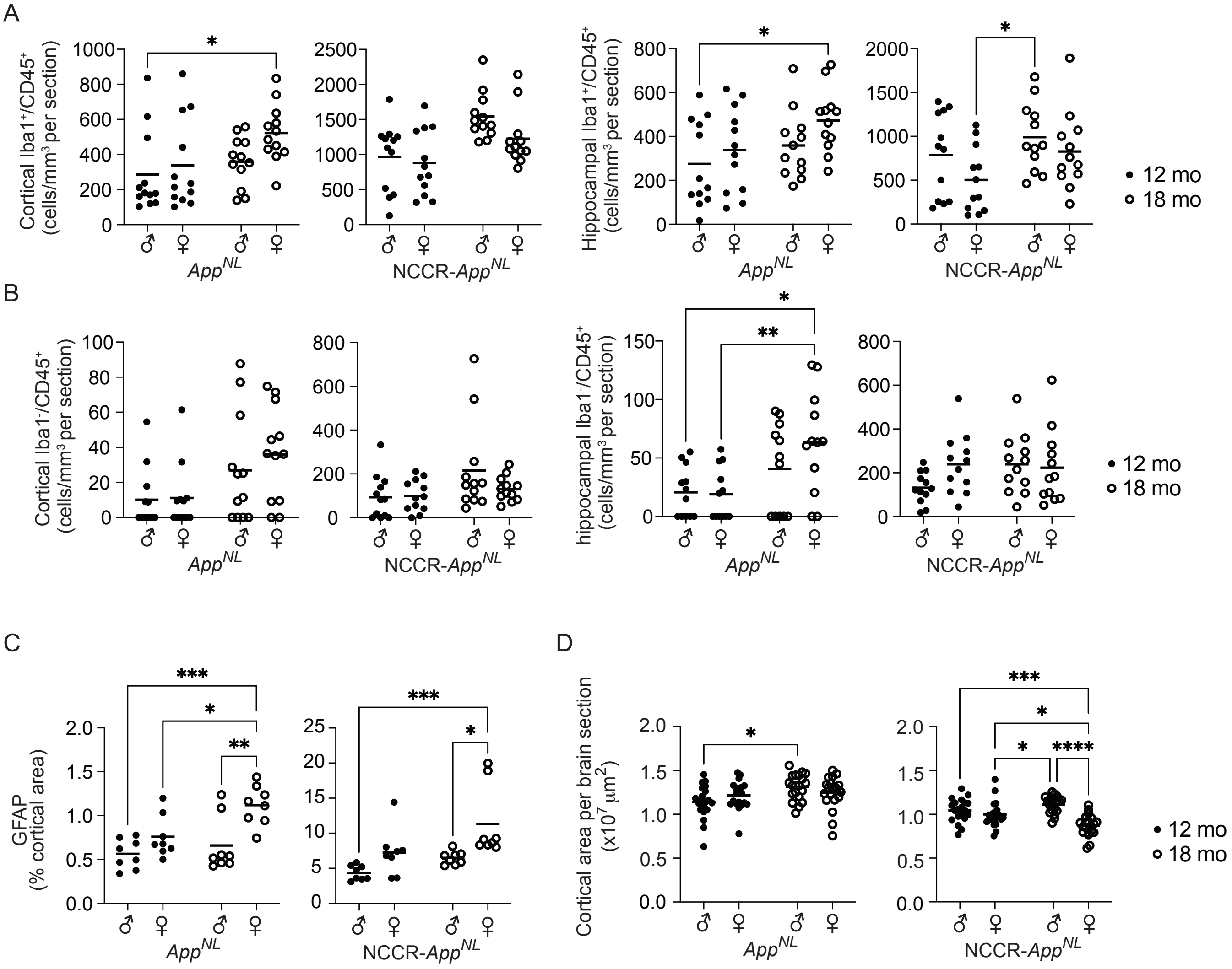

Supplementary figure 4

A

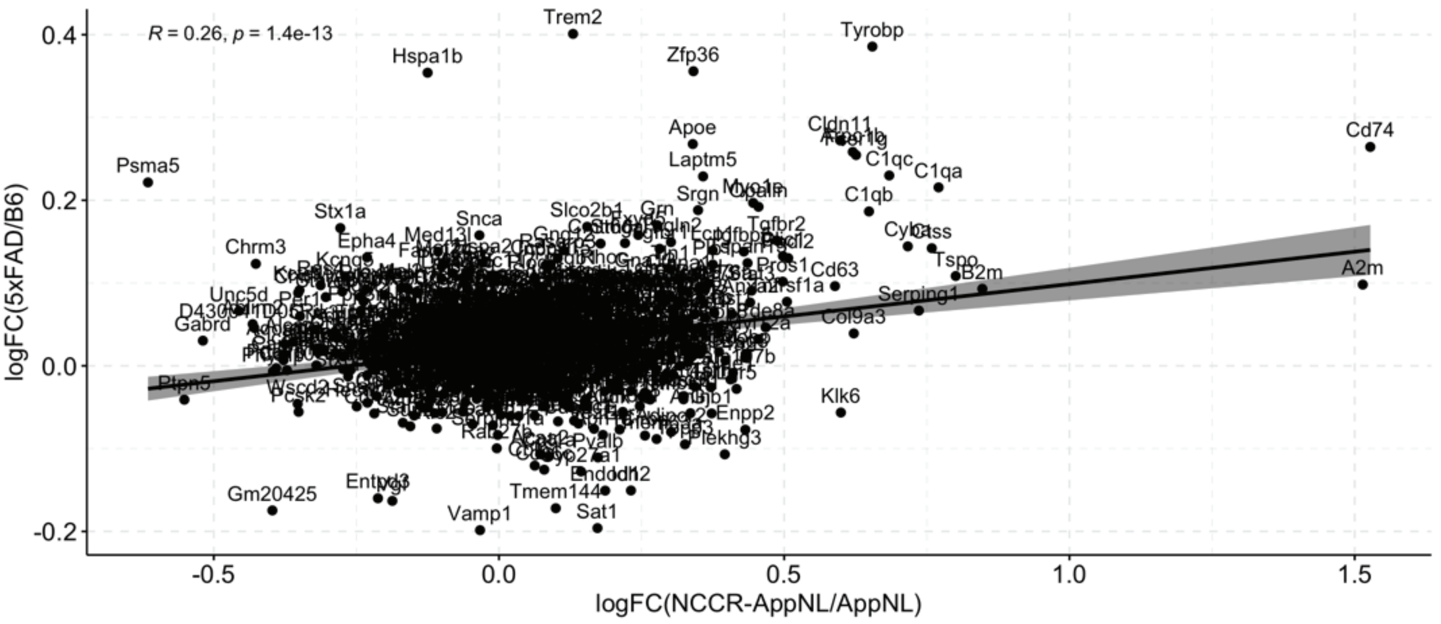


B

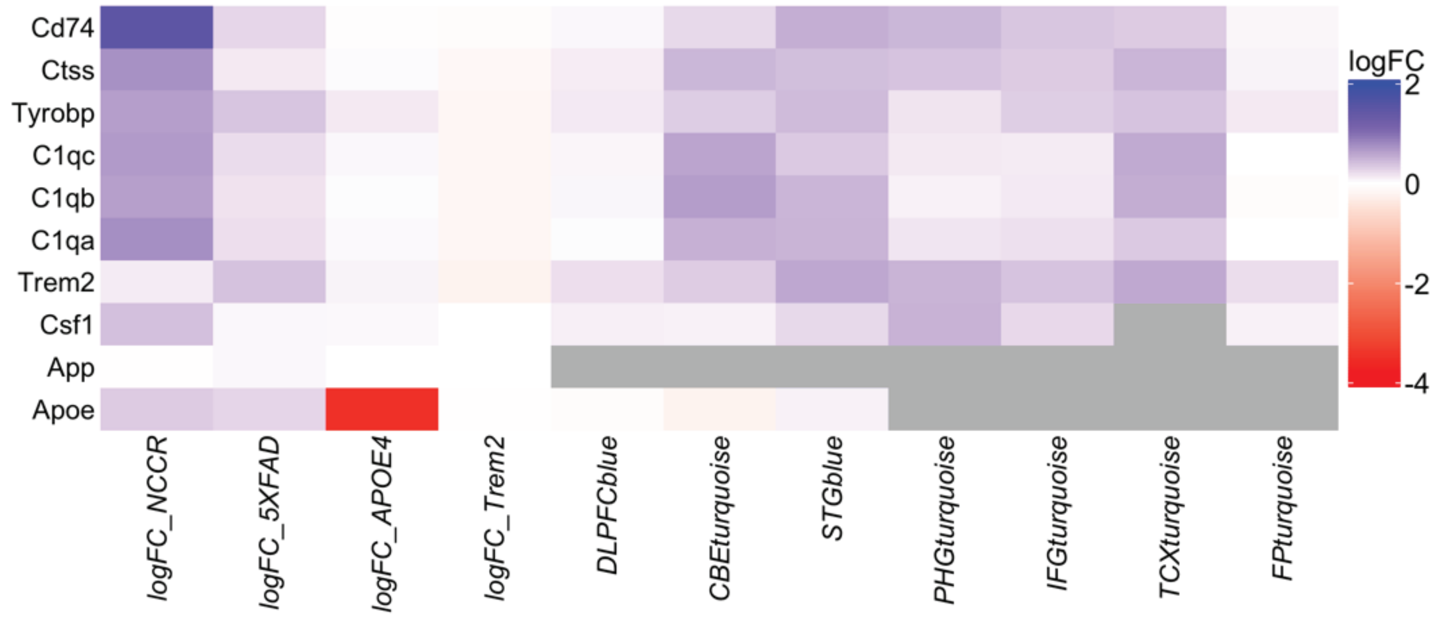

### Supplementary Figure 5

A

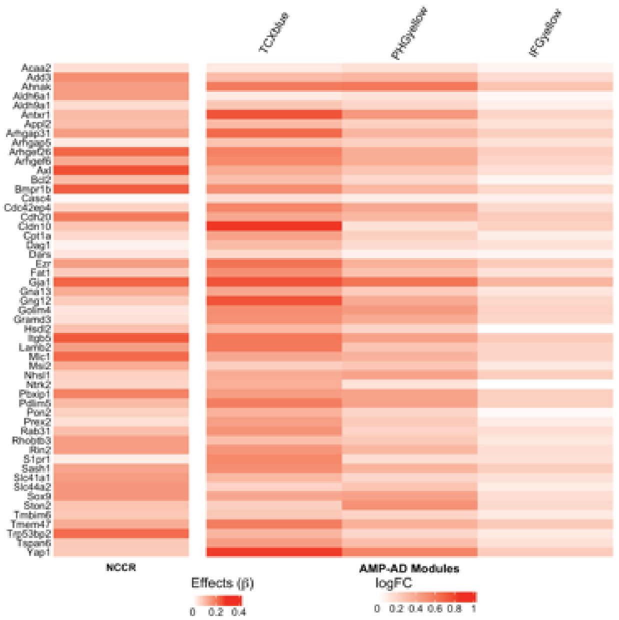


B

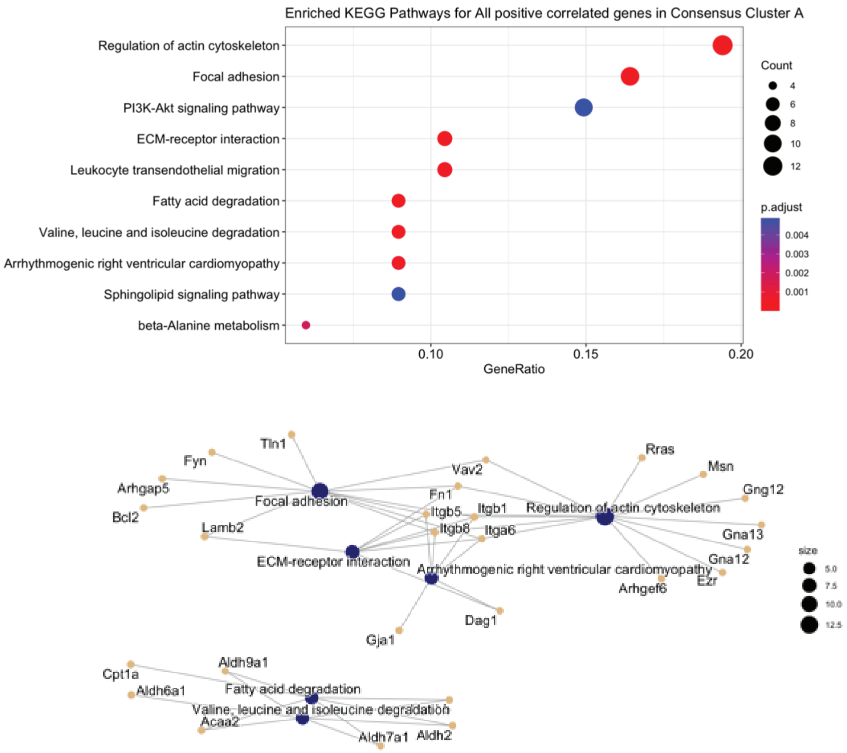

Suppl. Figure 6

A

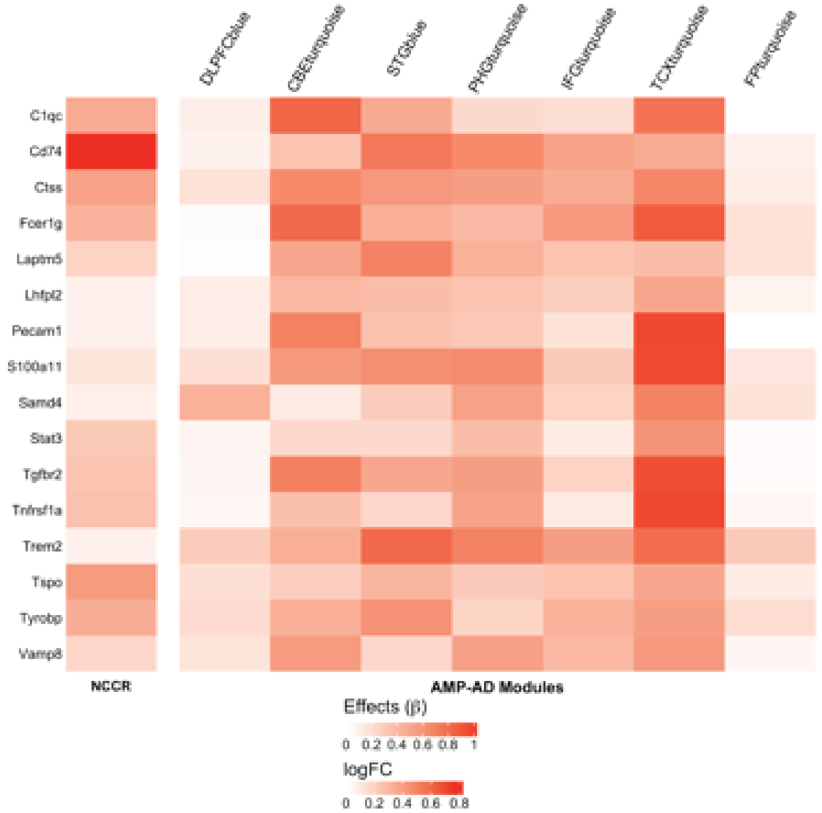


B

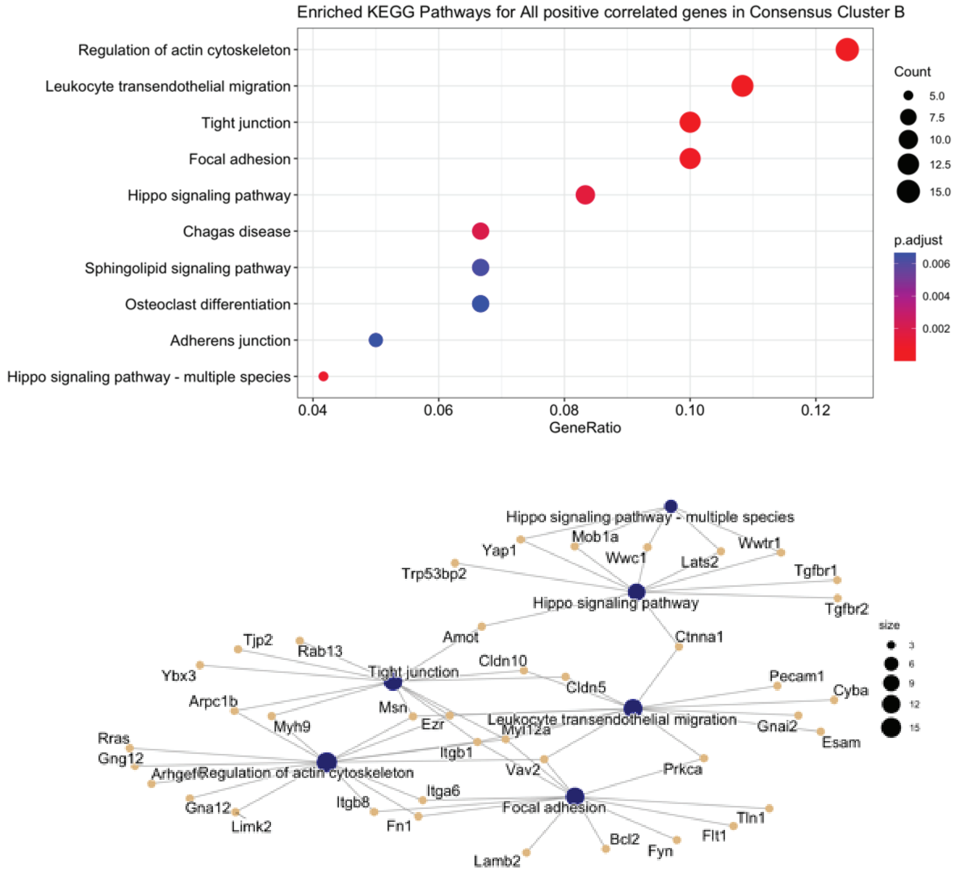

Suppl. Figure 7

A

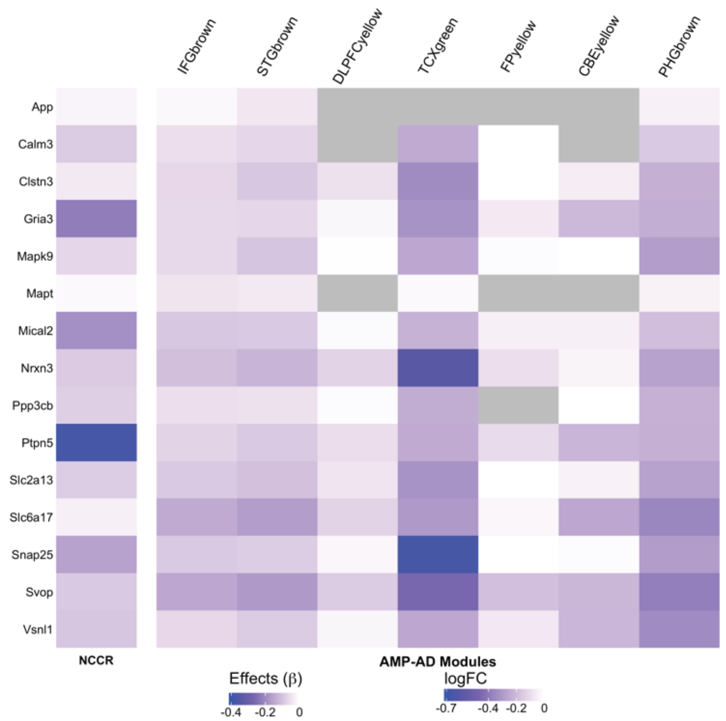


B

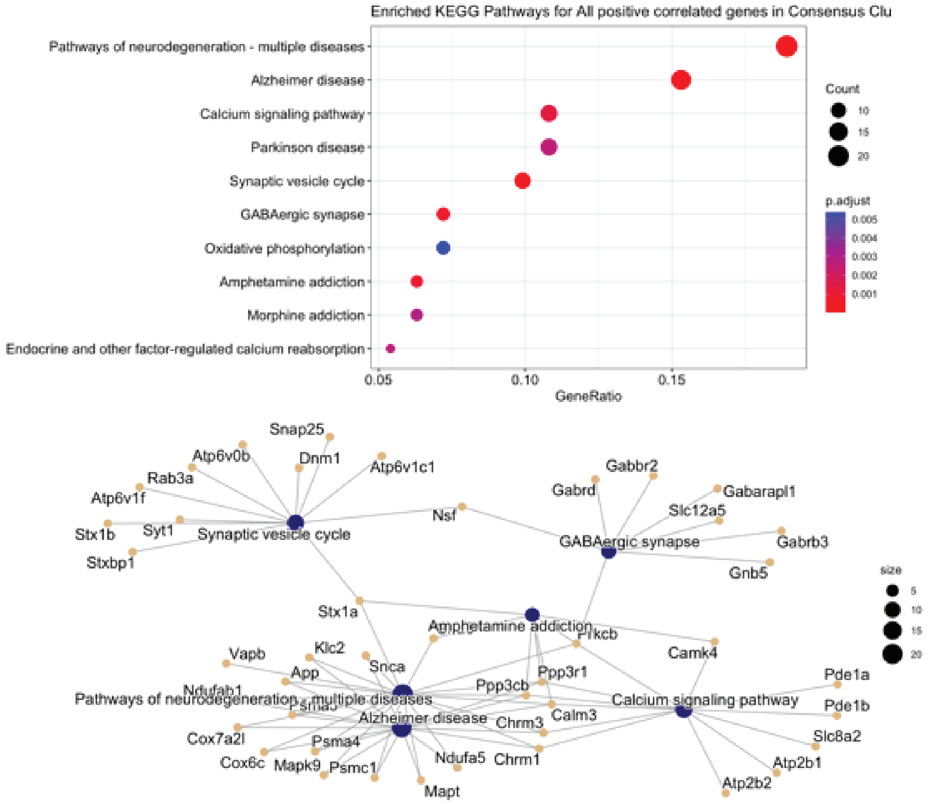

Suppl. Figure 8

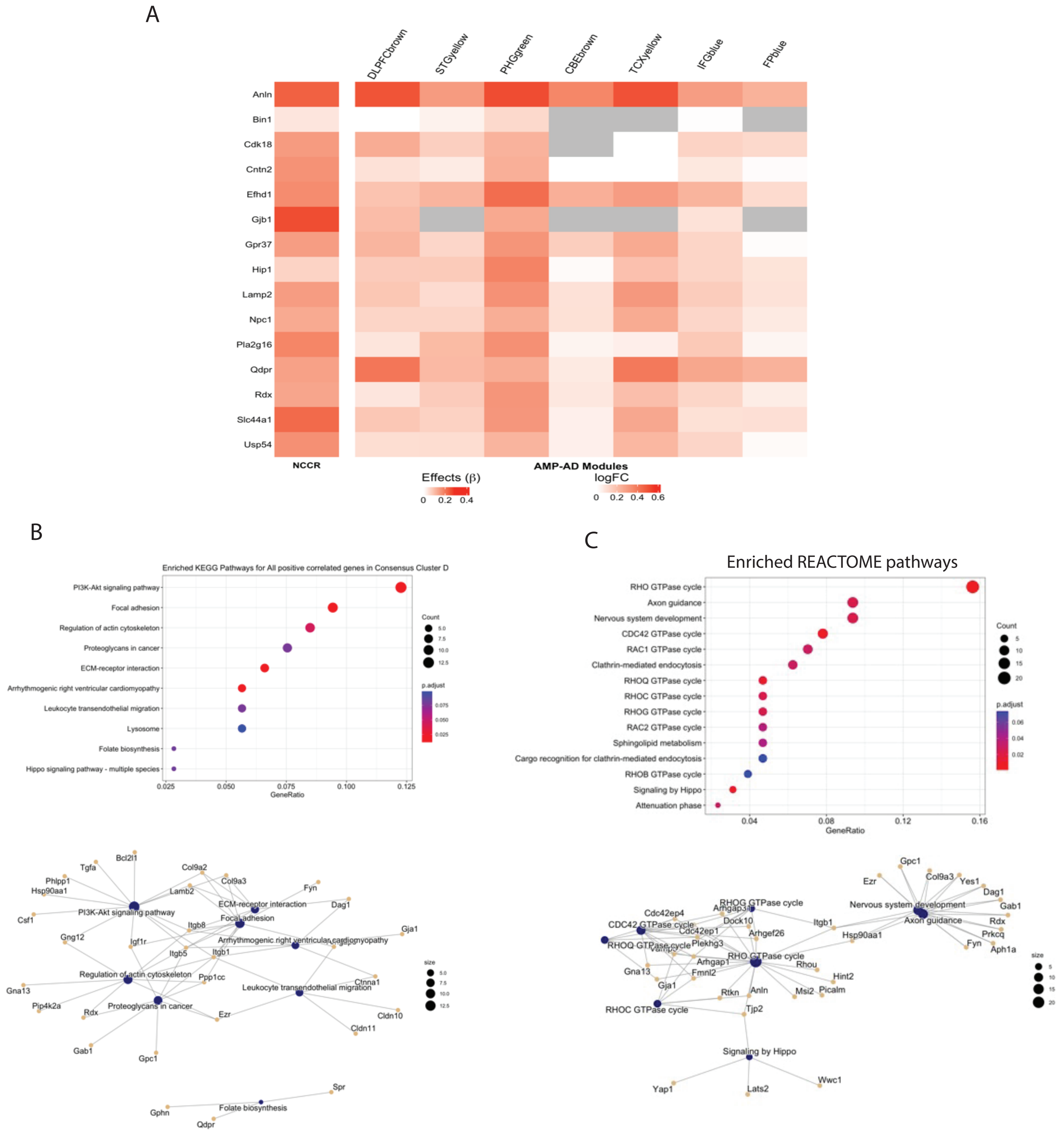

Supplementary Figure 9

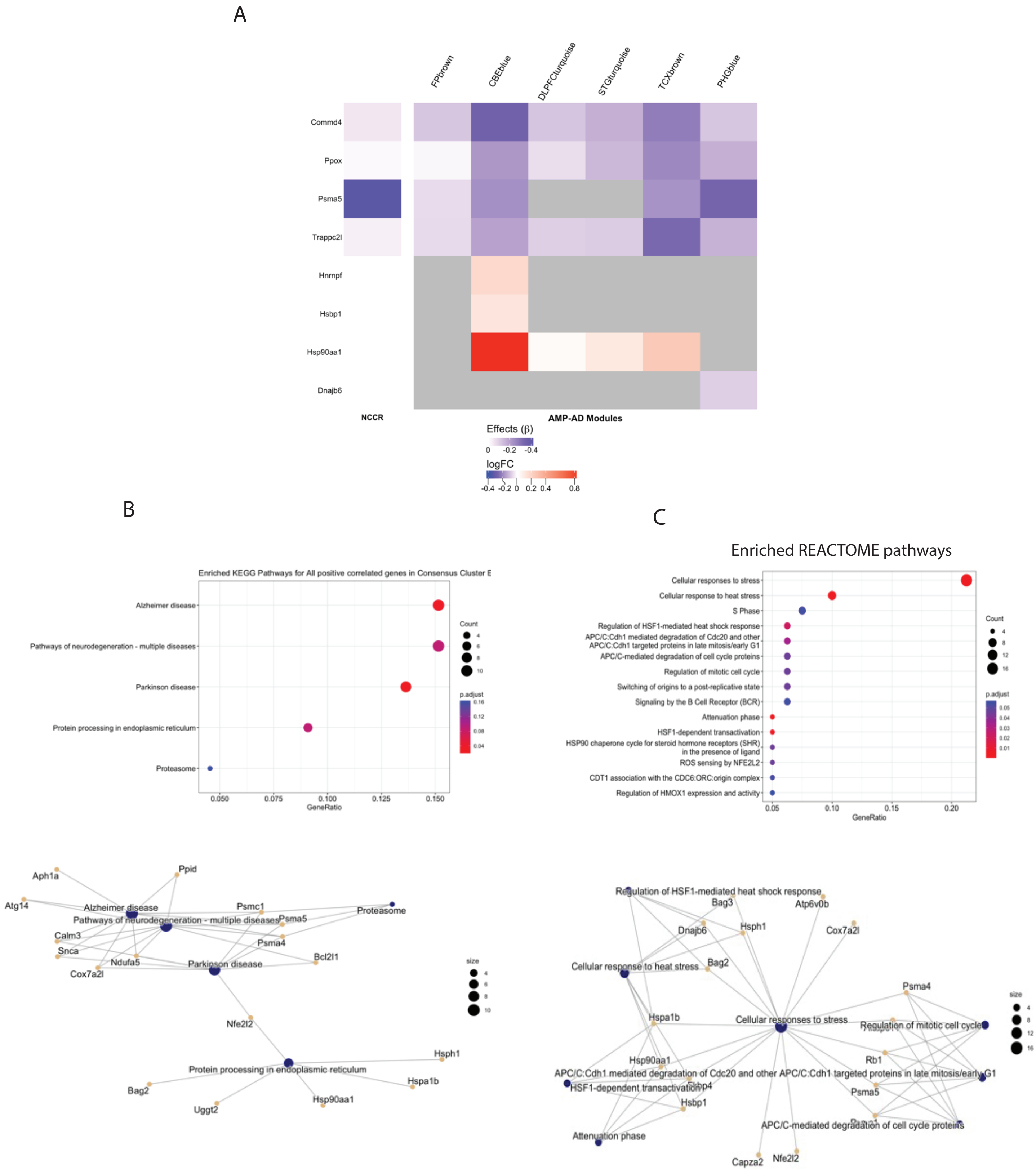

Supplementary Figure 10
LCM data results

A

#### SV40T-positive up-regulated genes

##### Enriched KEGG

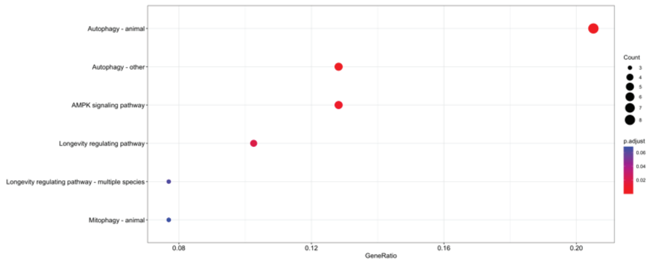


##### Enriched REACTOME

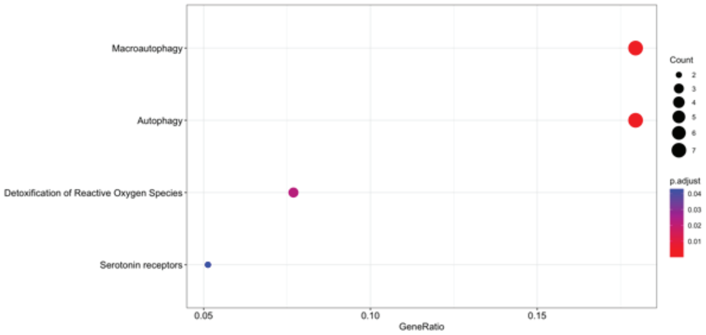


##### Enriched GO

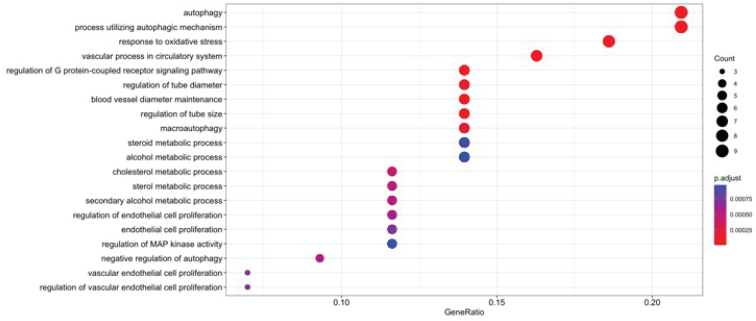


B

#### SV40T-positive down-regulated genes

##### Enriched KEGG

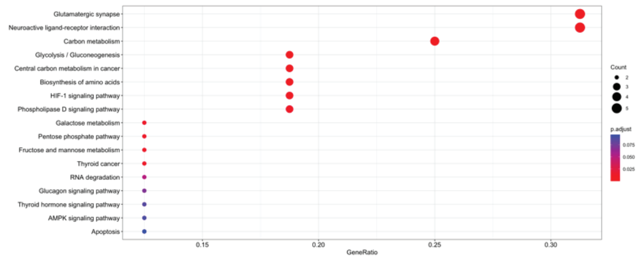


##### Enriched REACTOME

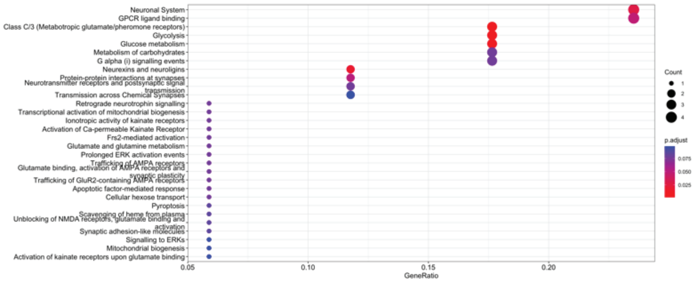


##### Enriched GO

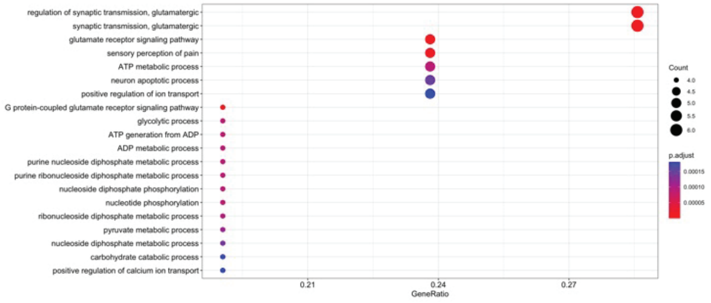
