## supplemental figure legends for "Aberrant neuronal cell cycle re-entry induces late-onset Alzheimer’s disease relevant neuropathological and gene expression changes"

**Supplemental figure legend**

**Suppl. Figure 1**

(A) NeuN labeled coronal brain sections were either co-labeled with either C-terminal Aβ42 (C42) or Aβ40 (C40) antibody. Intraneuronal Aβ(x-40) was observed in both NCCR-*App^NL^* (n=4 mice/sex/age) and *App^NL^* (n=4 mice/sex/age) animals (arrowheads). Intraneuronal Aβ(x-42) was only observed in NCCR-*App^NL^* animals (arrowheads). (B) No sex difference was observed in the number of cells bearing intraneuronal Aβ(x-42) labeling. Therefore, data from males and females were pooled for genotype x age two-way ANOVA analysis. (C) On the other hand, the number of Aβ(x-40) immunolabeled cell counts showed main effect difference for sex in NCCR-*App^NL^* animals but not for *App^NL^* animals (Supplementary Table 1). However, given that the post-hoc analysis did not reveal differences across NCCR-*App^NL^* samples, data from both males and females were pooled for genotype x age two-way ANOVA analysis. (D) 6E10 immunolabeling of coronal brain sections did not show Aβ plaques in either 12- or 18-month old *App^NL^* or NCCR-*App^NL^* mice. (E) Immunolabeling of coronal brain sections using PHF-1 antibody did not show PHF-1 phospho-tau lesions in either 12- or 18-month old *App^NL^* or NCCR-*App^NL^* mice. The numbers of matched coronal sections measured across the animals for the following analyses: Aβ(x-42), n=3; Aβ(x-40), n=1. Each data point represents a measure from each section.

**Suppl. Figure 2**

(A) No difference in main effect for sex was observed in the proportion of PCNA+ cells showing intraneuronal Aβ(x-42) co-labeling. Therefore, data from males and females were pooled for genotype x age two-way ANOVA analysis. (B) PCNA+ cells were similar between 12- and 18-month old NCCR-*App^NL^* animals, indicating that PCNA+ cells remains constant with age. Additionally, 18-month old animals put back on dox did not shown any PCNA+ cells demonstrating successful halting of NCCR using the dox diet (C) Hoechst signal intensity was measured for SV40T+ and PCNA+ cells and compared to SV40T- and PCNA- cells, respectively, within NCCR-*App^NL^* coronal brain sections as an indirect measure of DNA content. Post-hoc analysis showed statistically significant increase in the Hoechst signal intensity at 18-month old NCCR-*App^NL^* sample compared to all other groups (Tukey’s post hoc, p<0.0001). Following numbers of animals were evaluated: Regular diet groups: n=4 mice/sex/age/genotype. 18-month old back-on-dox groups: n=5 males for *App^NL^* and NCCR-*App^NL^* mice. The numbers of matched coronal sections measured across the animals for the following analyses: PCNA, n=1; Hoechst, n=2. Each data point represents a measure from each section.

**Suppl. Figure 3**

Sex effect was evaluated for neuroinflammation and cortical atrophy. No sex difference was observed­ for either(A) activated microglia or (B) brain leukocyte infiltration in the cortical and hippocampal areas. Therefore, data from males and females were combined for genotype x age two-way ANOVA analysis. On the other hand, sex difference was observed for (C) cortical area covered by GFAP signal and (D) cortical area measures. Therefore, data from males and females were separately analyzed in the main figures. Numbers of animals evaluated: n=4 animals/sex/age/genotype. The numbers of matched coronal sections measured across the animals for the following analyses: Iba1/CD45, n=3; GFAP, n=2; cortical area, n=5. Each data point represents a measure from each section.

**Suppl. Figure 4**

Gene expression changes in NCCR mice and 5xFAD mice were directly compared. (A) Correlation of expression changes in NanoString panel genes observed in NCCR-*App^NL^* mice compared to *App^NL^* mice with expression changes of these genes in 5xFAD mice compared to wild type B6 mice demonstrates significant positive correlation (R=0.26, p=1.4x10^-13^). Many of the neuroinflammation related genes show greater expression with NCCR compared to 5xFAD transgene expression. (B) Multiple AD risk genes are upregulated in both NCCR and 5xFAD mice and in immune system-related human co-expression modules. NCCR mice demonstrates the most robust upregulation of these genes, whereas these changes were unaffected in APOE4 and Trem2 LOAD risk gene mouse models.

**Suppl. Figure 5**

Pathway evaluation of genes from NCCR effect showing directional coherence with ECM-associated genes from human co-expression modules in Consensus Cluster A. (A) Heat map showing directional coherence between a subset of 128 NanoString panel genes for NCCR effect and AMP-AD human co-expression modules. (B) KEGG pathway enrichment analysis for 128 genes showing directional coherence s in Consensus Cluster A.

**Suppl. Figure 6**

Pathway evaluation of genes from NCCR effect showing directional coherence with immune system-associated genes from human co-expression modules in Consensus Cluster B. (A) Heat map showing directional coherence between a subset of 201 NanoString panel genes for NCCR effect and AMP-AD human co-expression modules. (B) KEGG pathway enrichment analysis for 201 genes showing directional coherence in Consensus Cluster B.

**Suppl. Figure 7**

Pathway evaluation of genes from NCCR effect showing directional coherence with neuronal system-associated genes from human co-expression modules in Consensus Cluster C. (A) Heat map showing directional coherence between a subset of 222 NanoString panel genes for NCCR effect and AMP-AD human co-expression modules. (B) KEGG pathway enrichment analysis for 222 genes showing directional coherence in Consensus Cluster C.

**Suppl. Figure 8**

Pathway evaluation of genes from NCCR effect showing directional coherence with cell cycle-associated genes from human co-expression modules in Consensus Cluster D. (A) Heat map showing directional coherence between a subset of 247 NanoString panel genes for NCCR effect and AMP-AD human co-expression modules. (B) KEGG pathway enrichment analysis for 222 genes showing directional coherence in Consensus Cluster D.

**Suppl. Figure 9**

Pathway evaluation of genes from NCCR effect showing directional coherence with organelle biogenesis and proteostasis-associated genes from human co-expression modules in Consensus Cluster E. (A) Heat map showing directional coherence between a subset of 141 NanoString panel genes for NCCR effect and AMP-AD human co-expression modules. (B) KEGG pathway enrichment analysis for 141 genes showing directional coherence in Consensus Cluster E.

**Suppl. Figure 10**

Pathway evaluation of 73 differentially expressed genes (fold change ≥ |2|) identified using laser capture microdissection microarray analysis of SV40T-immunopositive neurons compared to SV40T-immunonegative neurons from 4-month old NCCR mice. (A) KEGG and REACTOME pathway analysis for 47 upregulated genes. (B) KEGG and REACTOME pathway analysis of 26 downregulated genes.
